# Single-cell transcriptomic and epigenomic analysis reveals X-linked sex differences in aging mouse hypothalamus

**DOI:** 10.64898/2026.09.08.750001

**Authors:** Doudou Yu, Ilya Osipov, Lexi-Amber Hassell, Neta A. Shwartz, Kaitlyn H. Hajdarovic, Harold Marin, Ashley E. Webb

## Abstract

Sex differences contribute to brain aging, neurodegenerative diseases, and more broadly in determining rates of aging across species. The hypothalamus plays a central role in physiological homeostasis and healthy aging, yet how its cellular and molecular landscape diverges between males and females over the lifespan remains poorly understood. Here, we present a single-nucleus multi-omics analysis of the hypothalamus in young, middle aged, and aged male and female mice. We identified major hypothalamic cell types and characterized their sex- and age-dependent transcriptional and chromatin accessibility profiles. Notably, female-specific changes on the X chromosome (chrX) emerged as a prominent feature of aging, including changes to the X inactivation center and an overall increase in chrX gene expression and accessibility in immune cells and neurons. Pseudotime analysis of immune cells revealed an aging trajectory with sex-specific multi-omic programs, featuring increased inflammation in females compared to males. Delving deeper into the epigenetic signatures associated with these sex differences, we found that H3K27me3 – the repressive histone mark enriched on the inactive X in females – increased in abundance and underwent substantial genome-wide redistribution with age in both sexes, particularly on the inactive chrX in females. Collectively, these findings highlight distinct cell-type-specific aging trajectories in the male and female hypothalamus, identify female aging signatures associated with X-linked epigenetic regulatory programs, and provide a comprehensive resource for understanding the molecular basis of sex differences in brain aging.

## Introduction

Aging is a complex process characterized by progressive loss of cellular function across multiple organ systems. In the central nervous system, the hypothalamus plays a pivotal role in regulating metabolism, endocrine function, and circadian rhythms, positioning it as a key orchestrator of systemic health and aging^1–3^. Hypothalamic dysfunction has been linked to a number of physiological changes, including metabolic decline, altered sleep, and cognitive impairments in aging, yet how its cellular and molecular landscape changes with age in a sex-dependent manner remains poorly understood^1,4–6^. Because biological sex differences in lifespan, neurodegenerative disease risk, and immune function are well-established, dissecting the molecular underpinnings of sex-specific hypothalamic aging is critical for understanding the biology of aging and age-related disorders^7–9^.

Recent evidence implicates the X chromosome (chrX) as a potential determinant of sex differences in aging^10,11^. During mammalian development, one of the two X chromosomes in XX females is silenced to balance X-linked gene dosage with XY males, a process known as X chromosome inactivation (XCI)^12^. *Xist*, an X-linked long non-coding RNA that mediates XCI, increases with age in female neurons and may serve as a marker for neuronal aging^13^. Moreover, escape from XCI, in which certain genes evade transcriptional silencing, has been implicated in immune function, inflammation, and susceptibility to age-related disorders^14–16^. These observations suggest that chrX-linked changes may contribute to sex differences in brain aging, but the underlying mechanisms, key cell types, and regulatory dynamics are not known. XCI begins with coating of the future inactive X (Xi) *in cis* by *Xist*^17,18^, which directly or indirectly recruits various factors to facilitate silencing and chromatin remodeling^19–21^.

However, recent reports have identified widespread de-repression of previously silent genes on the Xi during aging (“age-related escape”) across multiple tissues, including liver, kidney, lung, heart, and brain, with the hippocampus examined specifically^22,23^; an allele-specific analysis of human RNA-seq data has since extended these observations of age-related loss of Xi silencing to humans^24^. While these age-associated chrX changes have been characterized at the transcriptomic level, whether they are associated with epigenetic remodeling of the X during aging remains unknown.

Epigenetic changes are a key hallmark of aging, spanning changes in individual chromatin marks to higher level genome organization. These include changes in DNA methylation, altered histone mark abundance and distribution, histone variant replacement, loss of heterochromatin, alterations in the activity of chromatin modifying complexes such as Polycomb Repressive Complex 2 (PRC2), and disruption to nuclear architecture, including erosion of lamina-associated domains (LADs) and topologically associating domain (TAD) boundaries^25–29^. Some of these changes can be reversed in certain contexts, such as cellular reprogramming, which has been interpreted as cellular rejuvenation^30^ and a path to future interventions.

Histone H3 lysine 27 trimethylation (H3K27me3) is a repressive histone mark deposited by PRC2 which plays a key role in epigenetic regulation of development, maintenance of cell identity, mammalian XCI, disease states such as cancer, and other contexts^31–33^. Notably, age-related changes in H3K27me3 distribution and total abundance have been reported in several tissues and cell types. Yang et al. observed an increase in total H3K27me3 abundance with age in mouse liver, and identified megabase-scale regions across the genome enriched for accrual of H3K27me3, which they termed “age domains”^34^. A similar spatial broadening occurs in mouse hematopoietic stem cells, where peak counts are comparable between young and aged cells yet H3K27me3 coverage expands with age^35^, and elevated H3K27me3 has also been observed in aged muscle stem cells^36^. Together, these studies indicate that H3K27me3 is redistributed, and in some tissues increased, during aging, raising the question of how these dynamics unfold on the X chromosome, where H3K27me3 has a specialized silencing role.

Advances in single-cell and single-nucleus ‘omics have enabled the high-resolution profiling of gene expression and chromatin accessibility across diverse cell types, providing powerful tools for dissecting aging trajectories in complex tissues such as the hypothalamus^37–41^. Prior studies have shown that age-associated transcriptional changes vary widely across hypothalamic cell types, with neurons, astrocytes, microglia, and oligodendrocytes exhibiting distinct aging signatures^13,38^. However, the extent to which these changes are sex-dependent, particularly at the epigenetic level, remains unclear. To address these gaps in knowledge, we performed single-nucleus (sn) multi-omic profiling of hypothalamic nuclei isolated from male and female mice at 3, 12, and 24 months of age, co-assaying snRNA-seq and snATAC-seq in the same nuclei to directly link gene expression with chromatin accessibility in individual cells. By leveraging HypoMap, a curated snRNA-seq reference atlas of the mouse hypothalamus^42^, we systematically annotated 13 major cell types from the transcriptomic modality and transferred these annotations to the epigenomic data. Our analysis revealed robust sex- and age-dependent patterns in chrX gene expression and chromatin accessibility, particularly in neurons, astrocytes, and immune cells. In female but not male mice, *Xist* expression and chrX accessibility increased significantly with age. To further investigate the epigenetic changes associated with chrX alterations, we performed genome-wide profiling of the histone mark H3K27me3 in young and aged hypothalami. We observed increased levels and substantial genome-wide redistributions of this histone mark with age in both sexes. Strikingly, we observed a widespread increase of H3K27me3 on the inactive X chromosome in aged female neurons that was not observed in males. Together, these findings provide new insights into the cell-type-specific molecular basis of sex differences in hypothalamic aging and highlight chrX changes as a key feature of female-biased aging trajectories.

## Results

### Single-nucleus multi-omics profiling of sex differences in the aging mouse hypothalamus

To investigate sex-specific aging dynamics in transcription and chromatin accessibility, we performed single-nucleus (sn) multi-omic profiling on nuclei isolated from male and female mouse hypothalami at 3, 12, and 24 months of age, pairing snRNA-seq with snATAC-seq (Assay for Transposase-Accessible Chromatin) on the 10x Genomics platform (Fig. 1a). Cell types were annotated by integrating snRNA-seq data with HypoMap, a curated mouse hypothalamus reference atlas^42^, and visualized in RNA-seq-only, ATAC-seq-only, and joint RNA-ATAC UMAP embeddings (Fig. 1b). After stringent quality control, 62,909 nuclei from 20 samples (n = 3–4 mice per age/sex group) were retained, yielding a dataset spanning 32,285 genes and 313,013 chromatin accessibility peaks (Fig. 1c, Extended Data Fig. 1). To validate cell identities, we generated two heatmaps, one of gene expression and one of gene activity derived from chromatin accessibility (ArchR^43^) for canonical marker genes across major hypothalamic cell types (Fig. 1d). Finally, we aggregated chromatin fragments by cell type using ArchR and applied MACS2^44^ to identify cell-type-specific chromatin accessibility peaks, which were visualized to confirm reproducibility (Fig. 1e).

**Figure 1.**
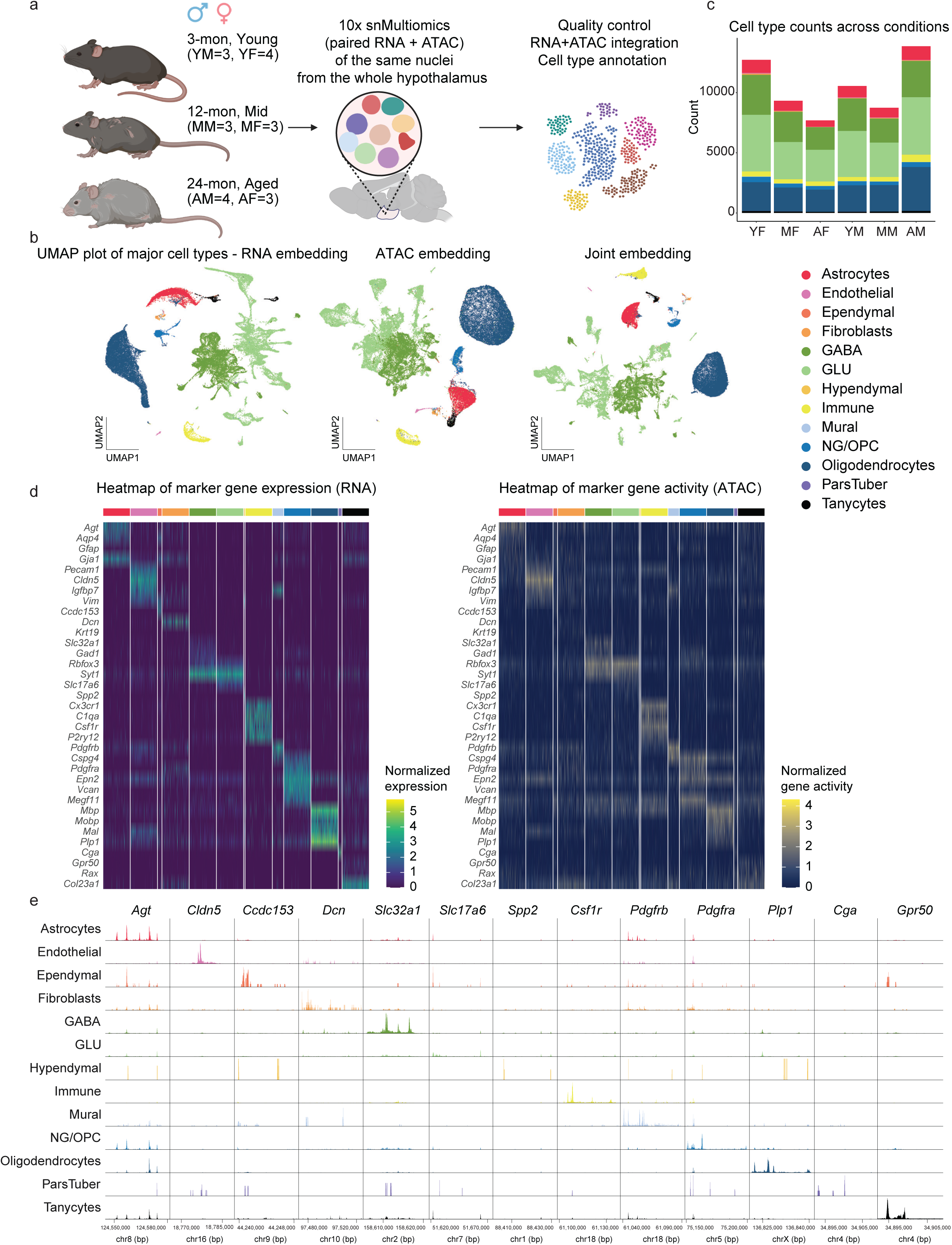
Single-nucleus multi-omics of the aging mouse hypothalamus. **a**, Experimental design: Single-nucleus RNA + ATAC profiling of hypothalamic cells from male and female mice at 3, 12, and 24 months. Nuclei were processed using 10x Multiome, followed by integration and cell type annotation. Group abbreviations for this and subsequent figures are YF: young female (n=4), MF: middle-aged female (n=3), AF: aged female (n=3), YM: young male (n=3), MM: middle-aged male (n=3), AM: aged male (n=4). **b**, UMAP of major cell types: RNA, ATAC, and joint embeddings showing distinct clustering of hypothalamic cell types. **c**, Cell type distribution: Bar plot showing nuclei counts across cell types, stratified by age and sex. **d**, Heatmaps of gene expression and chromatin accessibility: RNA (left) and ATAC (right) data highlighting marker gene expression and activity across major cell types. **e**, Pseudobulk Chromatin accessibility profiles: ATAC-seq signal at key marker genes across hypothalamic cell types.

### Sex-specific X-linked gene expression and chromatin accessibility changes across major cell types with age

Our previous work demonstrated that *Xist*, an X-linked long non-coding RNA essential for chrX inactivation^45^, is upregulated with age in female mice and may serve as a biomarker for neuronal aging^13^. To explore sex-specific dynamics, we analyzed age-related changes in *Xist* expression and chrX accessibility in both sexes. We observed a progressive increase in *Xist* expression and accessibility at the *Xist* transcription start site (TSS) in female mice across aging, with minimal expression and TSS accessibility in males, highlighting a robust female-specific expression pattern and increased accessibility with age (Fig. 2a-b, Extended Data Fig. 2a-b, Supplementary Table 1). These changes were most prominent in excitatory (GLU) and inhibitory (GABA) neurons, as well as immune cells. *Xist* expression was absent in young males, as expected, and males did not show an increase in *Xist* expression with age (Fig. 2a), indicating that the male-specific repression of *Xist* remains intact during aging in the hypothalamus.

**Figure 2.**
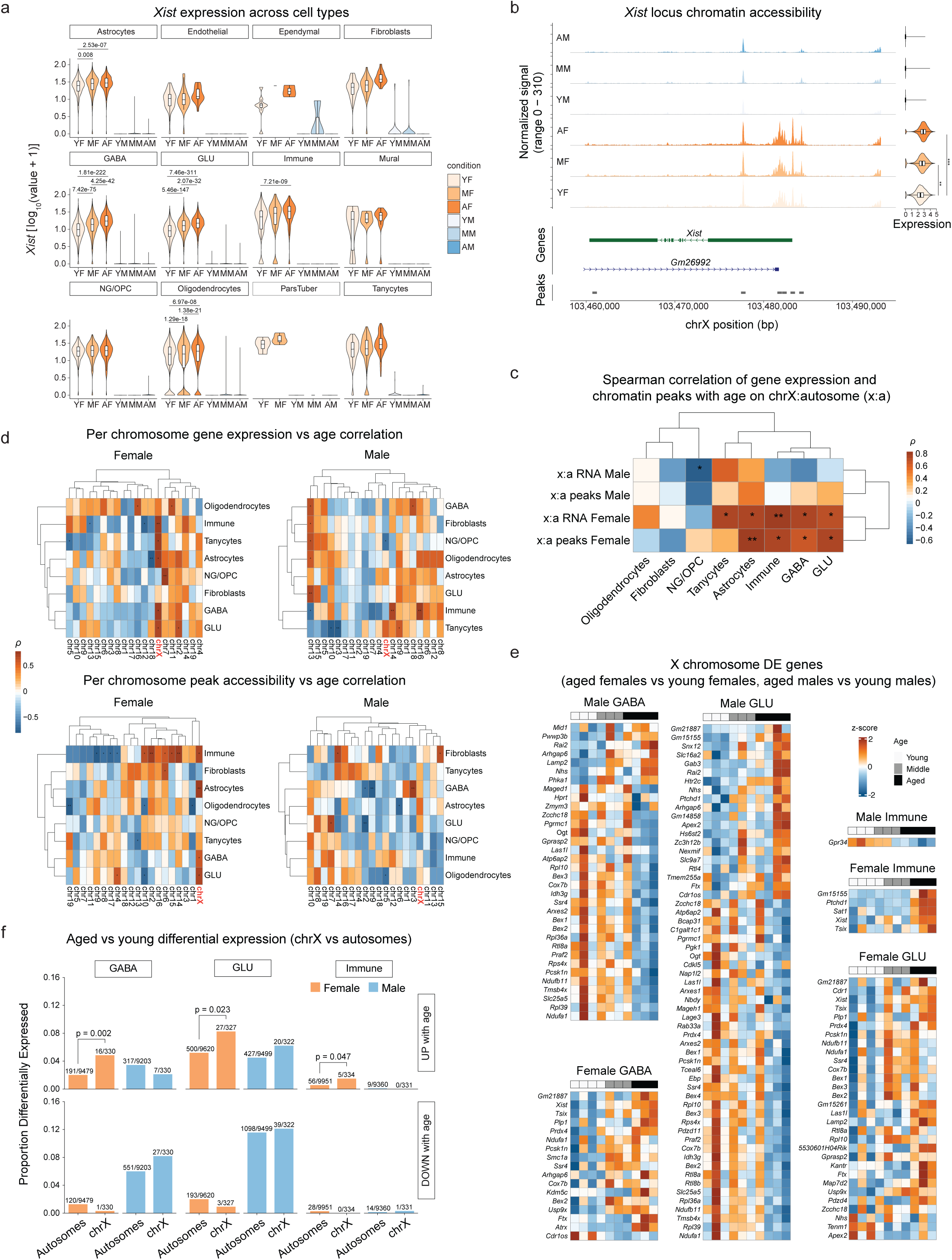
X-linked gene expression and chromatin accessibility changes across major cell types with age. **a**, Violin plots of *Xist* expression across cell types, stratified by sex, age, and condition. Bonferroni-adjusted p-values from the Wilcoxon test are shown above the violins for significant comparisons. **b**, Chromatin accessibility (left) and gene expression (right) at the *Xist* locus across groups for all cells, aggregated by age-sex combinations. Statistical significance for expression: Benjamini-Hochberg (BH) adjusted p-values from pseudobulk DESeq2 analysis; *adjusted p < 0.05, **adjusted p < 0.01, ***adjusted p < 0.001. **c**, Spearman correlations of gene expression and chromatin accessibility (chrX-to-autosome ratio) with age. Heatmap showing correlation patterns across major cell types. Significance: *p < 0.05, **p < 0.01. **d**, Heatmaps of Spearman correlations between chrX gene expression (top) and chromatin accessibility (bottom) in males and females with age. **e**, Heatmaps of significant aged vs young (Female: AF vs YF; Male: AM vs YM) chrX DE genes (p.adj < 0.05 and |log2FC| > 0.1) from the Wilcoxon test in three major cell types. Heatmap cells shows z-scored animal-level mean expression, with rows ordered by descending log2FC with age (from top to bottom). **f**, Proportion of expressed genes that were significantly differentially expressed on chrX vs autosomes, computed separately within each sex. Top row: significantly upregulated genes (p.adj < 0.05 and log2FC > 0.1). Bottom: significantly downregulated genes (p.adj < 0.05 and log2FC < -0.1). Fisher’s exact test nominal p-values shown above each comparison reaching p < 0.05.

We next assessed changes on the X chromosome relative to autosomes more holistically. Spearman correlation of the chrX-to-autosome (x:a) gene expression and chromatin accessibility ratios demonstrated a positive association between age and both x:a expression and accessibility in female GLU, GABA, and immune cells, but not in males (Fig. 2c). Analysis of individual chromosomes revealed that only chrX exhibited a positive age-related correlation in females (Fig. 2d, Extended Data Fig. 2b).

Chromatin accessibility analysis of the *Xist* promoter further revealed increased accessibility with age in females across major cell types (Extended Data Fig. 2c), reinforcing the sex and cell-type-specific epigenetic remodeling of chrX during hypothalamic aging. These findings highlight a sexually dimorphic regulatory landscape of the X chromosome during aging, in which *Xist* upregulation and increased chrX expression and accessibility mark female-specific aging trajectories.

### Transcriptional changes in major cell types reveal the X chromosome as a hub for sex differences in aging

Systematic analysis of differentially expressed genes (DEGs; identified by Wilcoxon rank-sum test, adjusted P < 0.05) across pairwise age comparisons revealed sex- and cell-type-specific aging signatures (Extended Data Fig. 1e, Supplementary Table 1).

Immune cells and astrocytes exhibited the most pronounced age-associated transcriptional changes in females, with the largest shifts occurring during the middle-aged to aged transition (immune cells, Aged female “AF” vs. Middle-aged female “MF”) and the greatest sex differences in aged populations (immune cells, AF vs. Aged male “AM”) (Extended Data Fig. 2c). In contrast, oligodendrocytes, GABA, and GLU neurons showed similar numbers of DEGs in both sexes, with few sex differences at mid-age. Notably, NG/OPC populations displayed the fewest DEGs with age, suggesting relative transcriptional stability (Extended Data Fig. 2d). Together, these findings highlight the sex-dependent dynamics of hypothalamic aging across cell types.

To dissect sex-specific aging signatures, we visualized within-sex pairwise DEG comparisons using scatterplots, plotting female log2 fold change (log2FC) against male log2FC, with DEGs color-coded as shared between sexes (purple), female-specific (red), or male-specific (blue) (Extended Data Fig. 2e). These comparisons revealed distinct temporal patterns: the early (mid-aged vs. young) and late-life (aged vs. mid-aged) transitions were dominated by female-biased changes (top and middle rows in Extended Data Fig. 2e), whereas the cumulative (aged vs. young) transition showed contributions from both sexes (bottom row in Extended Data Fig. 2e): astrocytes and immune cells remained female-biased, GABA and GLU neurons shifted toward male-biased changes, and oligodendrocytes exhibited balanced sex contributions. Fisher’s exact tests revealed significant enrichment of age-specific upregulated DEGs in female neurons for chrX genes, a pattern less pronounced in males (Fig. 2e–f), with 16/330 and 27/327 of expressed chrX genes being upregulated with age in GABA and GLU neurons, respectively. Interestingly, an overall greater number of downregulated genes was seen in males than in females between young and old GABA and GLU neurons (Fig. 2e-f, Extended Data Fig. 2e), with this pattern apparent both on autosomes and on chrX (e.g., of expressed autosomal genes, 551/9203 were downregulated with age in male GABA versus 120/9479 in female GABA). Collectively, these transcriptional analyses across major hypothalamic cell types converge on the X chromosome as a central hub of sex differences in aging, with female-biased, chrX-enriched changes distinguishing female from male aging trajectories.

### Female-biased disease-associated microglial signatures in the aging hypothalamus

Sex differences in the severity of inflammatory changes during brain aging have emerged in previous studies^46,47^. In aging and several neurodegenerative disease models, a subtype of activated microglia called “disease-associated microglia” (DAM) has been identified and found to have common transcriptional signatures across these age-related conditions. This transition begins with homeostatic microglia entering TREM2-independent DAM stage 1 (DAM1), which then facilitates entry into TREM2-dependent DAM stage 2 (DAM2), where upregulation of lysosomal, phagocytic, and lipid-processing genes, such as *Itgax* and *Lpl*, occurs^48–50^. The immune cell cluster in our dataset is composed predominantly of microglia and is hereafter referred to as microglia. Here, we observed an upregulation of DAM1 and DAM2 signatures in female microglia with age, consistent with our previous study^13^, whereas expression of DAM signatures appeared unchanged with age in the male hypothalamus (Fig. 3a).

**Figure 3.**
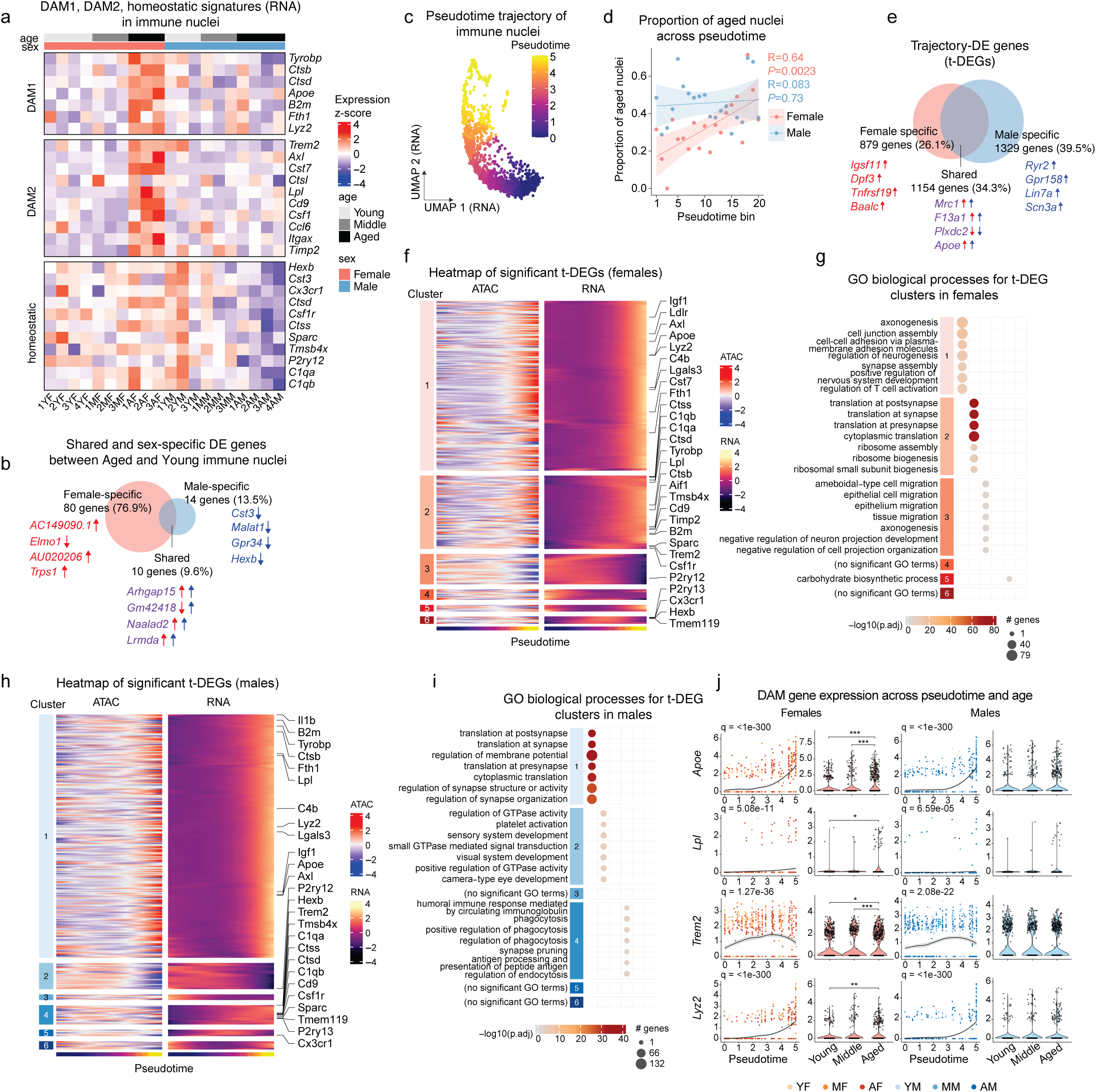
Female hypothalamic microglia show a stronger shift to a disease-associated state with age than males. **a,** Heatmap showing z-scores of pseudobulked expression in immune nuclei per animal for selected disease-associated and homeostatic microglial markers. **b,** Comparison of female (AF vs YF) and male (AM vs YM) aged-vs-young differentially expressed genes (p.adj < 0.05, Wilcoxon test) in immune nuclei. Genes listed are the top 4 most significant DE genes within each venn slice, with upward and downward arrows denoting upregulation and downregulation with age, respectively. Arrows are colored by sex (red: female; blue: male). **c,** RNA-derived UMAP embedding of the immune nuclei from the snMultiome dataset showing the pseudotime trajectory. **d,** Overlaid scatterplots showing, within each sex, the proportion of nuclei within each quantile bin that are aged. Immune nuclei were ordered by pseudotime and separated into 20 quantiles within each sex. Pearson correlations are shown. **e,** Comparison of female and male t-DEGs identified within each sex by Moran’s *I* test (Benjamini-Hochberg adjusted q-value (FDR) < 0.05). Top 4 significant t-DEGs within each venn slice are listed. Arrows: direction of expression changes across pseudotime with red arrows and blue arrows denoting females and males, respectively. **f,** Heatmap of significant t-DEGs in females. Columns are individual cells ordered by increasing pseudotime. Each gene’s normalized expression across pseudotime was smoothed using a spline model with 3 degrees of freedom and row-wise z-scored; rows were subsequently clustered hierarchically by Ward.D2 linkage using Pearson correlation distance and cut into 6 discrete groups. Accessibility was smoothed and z-scored in the same manner. **g,** GO biological process (BP) term over-representation analysis of the female t-DEG clusters from (f). Plotted are top terms with Benjamini-Hochberg adjusted p < 0.05. **h,** Same as (f) but showing male expression and accessibility of male t-DEGs. **i,** GO BP term over-representation analysis of the male t-DEG clusters from (h). Plotted are top terms with Benjamini-Hochberg adjusted p < 0.05. **j,** Normalized expression of individual DAM genes across pseudotime and age. Expression across nuclei is fitted by a locally estimated scatterplot smoothing (LOESS) curve (black line). Bonferroni-adjusted p-values from the Wilcoxon test are shown above each bracket.

Intriguingly, however, males showed a decrease in expression of homeostatic microglia markers with age that was not apparent in females (Fig 3a). More broadly, female microglia showed a higher number of differentially expressed genes from 3 months to 24 months of age compared to male microglia (Fig. 3b). Among the top male-specific age DE genes were *Cst3*, *Malat1*, *Gpr34* (an X-linked gene), and *Hexb*, all of which were downregulated with age and, apart from *Malat1*, are markers of homeostatic microglia.

To capture the continuum of gene expression states in our dataset’s immune nuclei, we performed trajectory inference^51^ across the transcriptional UMAP embedding of this population, after which each nucleus was assigned a pseudotime point based on its position along the trajectory (Fig. 3c). Aged female microglia were positioned later in pseudotime compared to young female microglia, with middle-aged female microglia occupying an intermediate pseudotime distribution (Extended Data Fig. 3a), indicating a progressive transition through this trajectory during aging. In contrast, male microglia did not appear to show a strong age-related shift toward later pseudotime (Extended Data Fig. 3a). When we divided microglia into equal-cell-number bins ranked by pseudotime within each sex, aged female microglia made up a progressively larger fraction of microglia in later pseudotime bins, whereas male microglia did not show this pattern (Fig. 3d). Similarly, the proportion of female microglia in each bin that were young significantly decreased along pseudotime, with an absence of this pattern in males (Extended Data Fig. 3b). This suggests that the inferred pseudotime trajectory captured microglial aging effects more strongly in females than in males. When evaluating gene expression and accessibility module scores for DAM signatures across the trajectory using AUCell^52^, we observed similar dynamics in both sexes: the DAM1 signature showed a progressive increase across pseudotime, whereas the DAM2 signature remained relatively stable, possibly owing to only a small subset of cells harboring non-zero expression of DAM2 genes (Extended Data Fig. 3d). Accessibility, on the other hand, was stable across the trajectory for DAM1 and DAM2 genes, while homeostatic genes underwent a loss of accessibility across the trajectory in both sexes (Extended Data Fig. 3d). Thus, the DAM and homeostatic gene sets are dynamic in both males and females across pseudotime, but the trajectory only captures aging in the female microglia.

To expand the analysis from curated DAM signatures toward broader transcriptional and chromatin accessibility changes, Moran’s *I* test was applied within each sex with an FDR cutoff of 0.05 to obtain sets of genes that were significantly differentially expressed and sets that were differentially accessible across the trajectory (Supplementary Table 2). Among the 3,362 trajectory differentially expressed genes (t-DEGs) identified, 34.3% were shared across sexes (Fig. 3e). We also observed a similar degree of overlap (24.6%) between sexes for genes that had differential accessibility (t-DA genes) across the trajectory (Extended Data Fig. 3c, Supplementary Table 2). We next visualized t-DEG dynamics across the trajectory and clustered them into discrete groups, finding that upregulated t-DEG clusters notably contained several DAM1 and DAM2 signatures, and that accessibility patterns appeared to generally track with gene expression (Fig. 3f, 3h). We performed GO term overrepresentation analysis and found that upregulated female t-DEG clusters were enriched for nervous system related Gene Ontology (GO) terms including axonogenesis, synapse assembly, and positive regulation of nervous system development (Fig. 3g, Supplementary Table 2). Upregulated male t-DEG clusters were enriched for phagocytosis, endocytosis, and synapse pruning terms (Fig. 3i, Supplementary Table 2). Both sexes showed increased expression of synapse and translation related GO terms across the trajectory, whereas males showed decreased expression of GTPase related GO terms (Fig. 3g, 3i, Supplementary Table 2). Similar patterns were seen in the accessibility modality, with males showing decreased accessibility in GTPase pathways, and both sexes showing declining accessibility of migration related pathways (Extended Data Fig. 3e-3h, Supplementary Table 2).

Notably, *Apoe*, a DAM marker and Alzheimer’s disease (AD) risk factor gene, was significantly upregulated with age in females, but not in males. Despite this, both sexes showed upregulated *Apoe* expression across the trajectory. This same pattern held true for other DAM genes including *Lpl*, *Trem2*, *Lyz2*, *Tyrobp*, and others (Fig. 3j, Extended Data Fig. 3i). Other markers such as *Cst7*, a DAM marker which has been shown to regulate microglia in a sex-dependent manner in a mouse model of AD^53^, and *Lgals3*, whose transcription is generally known to increase during aging^54^, showed stronger upward trends in female microglia with age than males (Extended data Fig. 3i). Despite males showing stronger downregulation of homeostatic signatures with age compared to females, the homeostatic marker *P2ry12* was downregulated with age in both sexes (Extended Data Fig. 3i). In female microglia, upregulation of individual DAM genes was conspicuously visible at the 24-month timepoint (Fig. 3j, Extended Data Fig. 3i), especially in the case of DAM2. Together with the pseudotime analysis, this suggests that the changes to microglial states begin early and strengthen between 12 and 24 months. Collectively, our data suggest a stronger age-related induction of a DAM-like state in female microglia than in males, with broader female-specific transcriptional shifts with age also being identified by pseudotime analysis. The overall sex-specific pattern we observed, where microglia shifted towards later pseudotime only in females (Fig. 3d), is intriguing given that both sexes shared similar dynamics across the trajectory itself. We interpret this as male and female microglia possessing a similar continuum of transcriptional profiles, but only the female population drifting toward the “activated” side of this continuum with age.

### H3K27me3 levels increase with age in the hypothalamus, with a female-specific increase and redistribution on the X chromosome

X chromosome silencing during development is initiated by coating of the chromosome in *cis* by *Xist*, followed by widespread deposition of repressive histone modifications including the heterochromatin mark H3K27me3. Increased total levels of H3K27me3 with age have been reported in mouse liver^34^ as well as in other species, including killifish brain^55^ and muscle^56^, *Drosophila* head and muscle^57^, and post-mortem brain from aged humans^58^. In contrast, a loss of H3K27me3 has been reported in aged mouse epidermis^59^, suggesting tissue-specific aging changes in this histone mark’s abundance. To investigate the extent to which the changes in *Xist*, global chrX gene expression, and chromatin accessibility are associated with heterochromatin changes, we profiled H3K27me3 in the aging hypothalamus using ChIP-seq on 3 month-old and 24 month-old mice of both sexes (Fig. 4a). To capture potential differences in the total levels of H3K27me3 between conditions, we incorporated spike-in normalization into our experimental design by adding a small, constant quantity of exogenous *Drosophila* chromatin to each sample prior to chromatin immunoprecipitation (ChIP-Rx)^60,61^.

**Figure 4.**
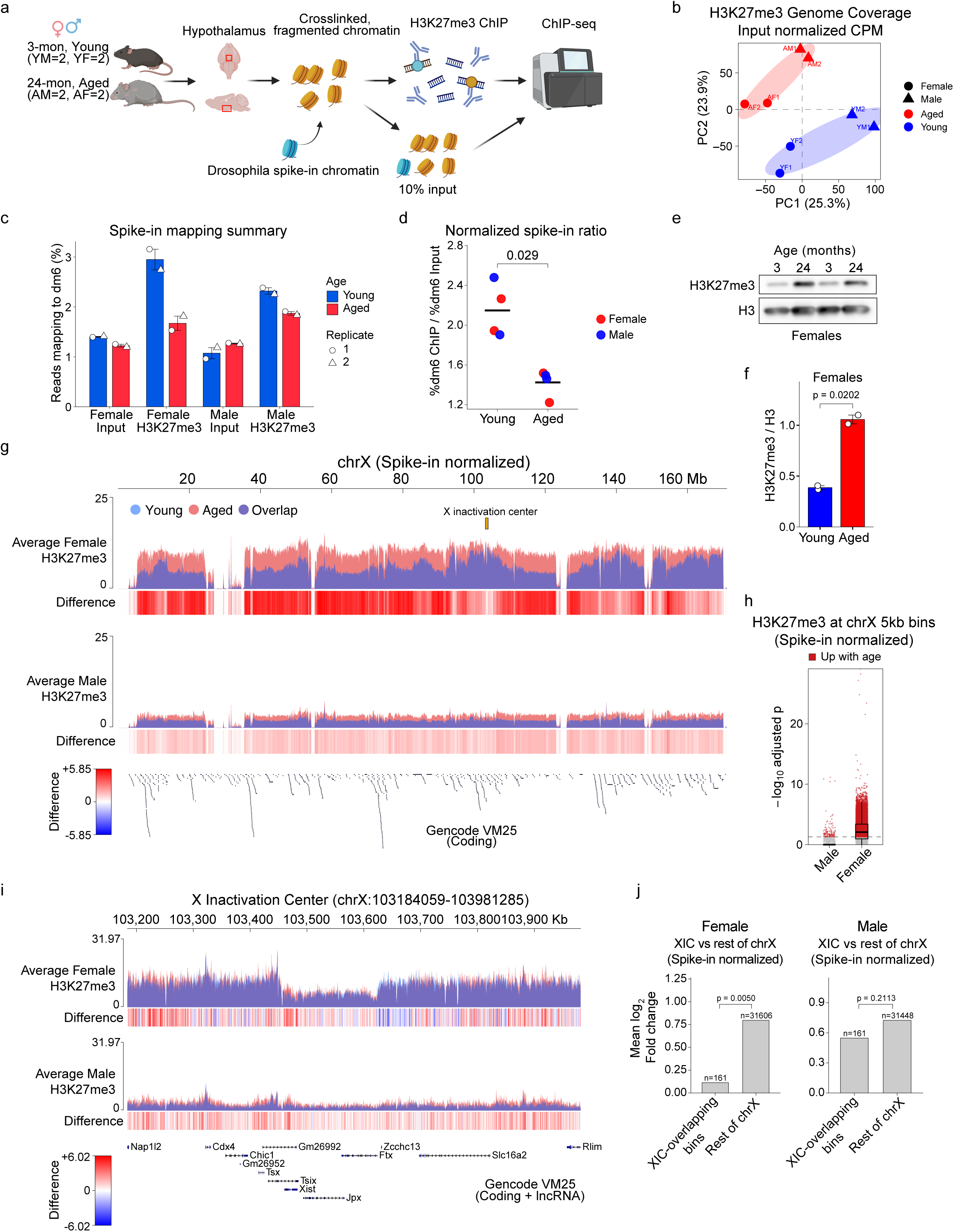
H3K27me3 levels increase and redistribute with age in the hypothalamus, with female-specific changes on the X chromosome. **a,** Experimental design for *Drosophila* spike-in normalized ChIP-seq of H3K27me3 in 3-month-old and 24-month-old mouse hypothalamus samples of both sexes (YM: young male (n=2), YF: young female (n=2), AM: aged male (n=2), AF: aged female (n=2)). **b,** Principal component analysis (PCA) of input-normalized H3K27me3 enrichment (5-kb genomic bins). **c,** Alignment rate to the *Drosophila* spike-in genome (dm6) during mapping of H3K27me3 ChIP-seq reads to the concatenated mouse-*Drosophila* (mm10-dm6) reference genome. **d,** Spike-in alignment rate in ChIP samples normalized to the spike-in alignment rate in matched input samples. Two-sided Wilcoxon rank-sum / Mann-Whitney U test, n=4 biological replicates per age group. Young and aged means are represented by horizontal black bars. **e,** H3K27me3 western blot of acid-extracted histones from hypothalamus samples of 3-month-old and 24-month-old female mice (n=2 animals per age group). A separate set of animals was used for western blotting. **f,** Quantification of western blot from (e). Two-sided unpaired Welch’s t-test with bars representing mean ± SEM **g,** X chromosome genome browser tracks of spike-in normalized H3K27me3 ChIP-seq signal. Within each sex, average aged signal (red) is overlaid with average young signal (blue). XIC: X Inactivation Center. **h,** H3K27me3 spike-in normalized differentially enriched regions (DESeq2) across 5-kb bins for the X chromosome (p.adj < 0.05 and |log2FC| >= 0.5). Non-significant bins are shown as grey dots. **i,** Genome browser tracks of spike-in normalized H3K27me3 ChIP-seq signal at the X inactivation center (chrX:103184059-103981285) for females and males. **j,** Mean log_2_ fold-change in H3K27me3 signal across 5-kb bins, where XIC-overlapping bins were defined by coordinates chrX:103184059-103981285 and compared to the remaining chrX bins. The number of bins (n) over which the mean log2fold change was taken is displayed above each bar. Empirical p-values are from an equal-width genomic window permutation test comparing changes with age at the XIC versus the X chromosome background (see Methods).

Principal component analysis of input-normalized H3K27me3 coverage showed clustering of samples by sex and age in the first two principal components, respectively, demonstrating a separable contribution of both sex and age to the variation in our ChIP-seq dataset (Fig. 4b). Globally, we observed that aged ChIP samples had a lower proportion of reads mapping to the *Drosophila* spike-in genome (dm6) than young samples, consistent with increased immunoprecipitation of mouse H3K27me3 in aged samples driving down their relative proportion of spike-in reads, particularly in females (Fig. 4c). To quantify this change and account for technical variability in the initial addition of spike-in chromatin, we normalized each ChIP sample’s spike-in mapping rate to that of its matched input and observed a significantly lower spike-in mapping rate with age, consistent with increased H3K27me3 abundance in the aged mouse hypothalamus (Fig. 4d). We confirmed this finding by western blot, detecting increased H3K27me3 in 24-month-old versus 3-month-old females in hypothalamus histone extracts isolated from an independent set of mice (Fig. 4e-f), and further validated with an additional set of 3-month-old and 21-month-old mice, again showing an increase in H3K27me3 with age in both sexes, with the increase appearing more pronounced in females than males (Extended Data Fig. 4a).

In examining the H3K27me3 distribution across the genome at higher resolution, we found – across multiple autosomes – the appearance of large megabase-scale regions that gained H3K27me3 with age in both sexes (Extended Data Fig. 4b). These regions resemble the H3K27me3 “age domains” which have previously been reported in aged mouse liver, kidney, muscle, and heart^34^. Interestingly, when integrating our H3K27me3 data with the snRNA-seq and snATAC-seq data, we observed that the hypothalamic age domains occurred in areas with low/near-absent accessibility and gene expression (Extended Data Fig. 4b; see shaded grey bar examples), suggesting these domains occur at silenced heterochromatin.

Considering the sex-specific role of H3K27me3 on the inactive X chromosome, we next examined whether age-related changes on the X chromosome differed between sexes. In contrast to autosomal changes (Extended Data Fig. 4b), we observed striking sex differences in H3K27me3 enrichment on the X during aging (Fig. 4g). Females showed a chromosome-wide gain of H3K27me3 across the X with age (Fig. 4g), with an appreciable feature being the flatter, more uniform H3K27me3 profile in aged females compared to the distinct “peaks and valleys” in young females (Fig 4g). This age-related flattening was more apparent when normalizing ChIP coverage to input only, and was not seen in males (Extended Data Fig. 4c). To obtain an unbiased quantitative assessment of H3K27me3 distribution changes during aging, we divided the genome into non-overlapping 5 kilobase (kb) bins and performed differential enrichment analysis using DESeq2^62^(Supplementary Table 3), incorporating spike-in normalization into the analysis (see Methods). Bins harboring differential H3K27me3 with age largely overlapped between males and females, though not fully, demonstrating shared and sex specific changes in this mark, with females having an overall higher number differentially bound bins (Extended Data Fig. 4d). This can be in part explained by the larger number of X chromosome bins that were significantly enriched for H3K27me3 increases with age in females compared to males (Fig. 4h), whereas autosomes were relatively similar in enrichment between males and females (Extended Data Fig. 4e).

Intriguingly, in females, the X inactivation center (XIC; chrX:103184059-103981285)^63^ gained less H3K27me3 with age than other regions of the X (Fig. 4g, 4i, 4j), representing a “cold spot” on the X that was less prone to age-related increases in this mark, which is consistent with a continued requirement of *Xist* transcription to maintain XCI. This observation is also consistent with female upregulation of XIC transcripts such as *Xist* and *Tsix* with age in the snRNA-seq and snATAC-seq data (Fig. 2b). Still, spike-in normalization suggests that absolute H3K27me3 levels are relatively stable in this region with age (Fig. 4i). We further found that gene poor regions had more pronounced gains of this mark compared to gene dense regions, with this observation also extending to the female X chromosome (Extended Data Fig. 4f). Additionally, input-normalized H3K27me3 coverage was negatively correlated with our pseudobulked snRNA-seq and snATAC-seq coverage, consistent with H3K27me3’s role in transcriptional repression and heterochromatinization (Extended Data Fig. 4g).

Collectively, our data show that aging is associated with a global increase in H3K27me3 in the hypothalamus, the appearance of autosomal age domains at heterochromatin across the genome, and pronounced female-specific elevation and flattening of H3K27me3 across the X chromosome, with the possible exception of the XIC and gene-dense regions, which appear relatively spared from H3K27me3 accrual.

### *Xist* RNA and H3K27me3 increase with age on the inactive X in female neurons

Our snMultiome, ChIP-seq and western blot data showed that *Xist* and H3K27me3 increase with age in the female hypothalamus, raising the question of whether these changes occur on the active or inactive X chromosome. To distinguish these possibilities we turned to imaging, which preserves spatial information, combining RNA fluorescence in situ hybridization (FISH) for *Xist* with H3K27me3 immunohistochemistry in hypothalamic neurons from young, middle-aged, and aged mice of both sexes (Fig 5a). Interestingly, aged female neurons showed a trend toward increased *Xist* area relative to nuclear area compared with young- and middle-aged neurons (Fig 5a and Extended Data Fig. 5a; p = 0.0546), and a significant increase in *Xist* intensity relative to young neurons (Extended Data Fig. 5b; p = 0.0101). No *Xist* signal was detected in male neurons at any age (Extended Data Fig. 5c), confirming both the specificity of probe hybridization in female mice and that the age-associated *Xist* increase is sex-specific. *Xist* RNA levels therefore not only intensify with age but also trend toward greater nuclear coverage, consistent with increased *Xist* RNA accumulation, and possibly spreading. Alternatively, this increase in relative area could reflect a spatial increase in Xi volume, marking a decompaction of the Barr body with age despite increasing *Xist* intensity.

**Figure 5:**
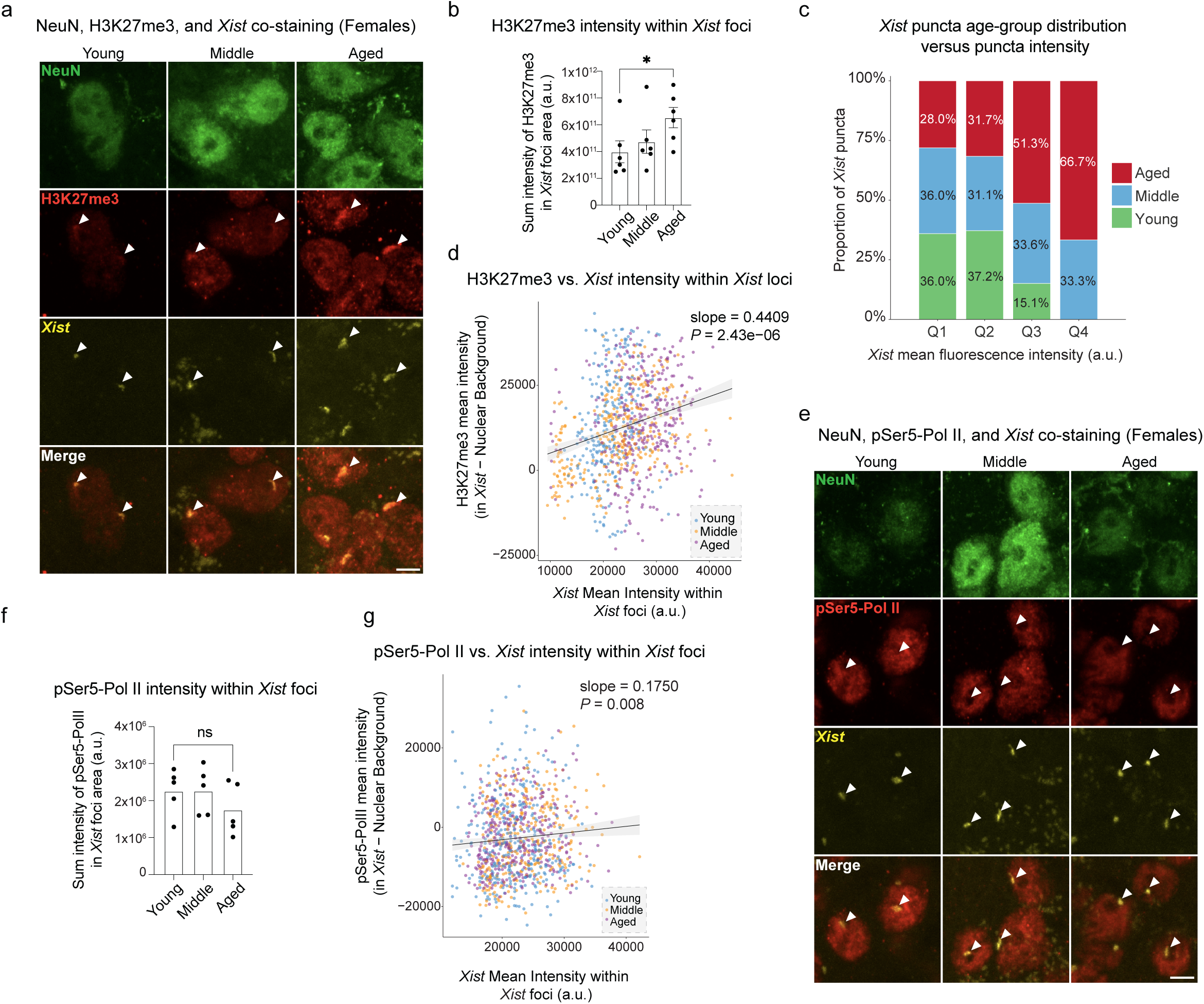
*Xist* RNA and H3K27me3 are enriched on the inactive X with age in neurons of the female hypothalamus. **a,** Representative confocal images of NeuN, *Xist*, and H3K27me3 in coronal sections of the arcuate nucleus in the hypothalamus from Young (4-month-old), Middle-aged (12-month-old), and Aged (22-month-old) female mice (same ages throughout the figure). Merged images show the colocalization of *Xist* and H3K27me3 within neuronal nuclei. Images were acquired using a 63x with oil immersion. Scale bar, 5 µm. Images are representative of n = 6 animals per age group. **b,** Quantification of H3K27me3 intensity within *Xist* foci across ages in neurons from female mice. Each point represents the mean across all neurons per animal, with bars indicating mean ± standard error of the mean (SEM) per age group. Statistical significance was assessed using unpaired t-test. n = 1,623 neurons from 18 animals imaged. **c,** Proportion of *Xist* RNA FISH foci from each age group within equal-width bins of *Xist* mean fluorescence intensity (bin width: 8,694 a.u.). Pooled from H3K27me3 and PolII imaging datasets (total n = 1,986 foci; 317, 1,175, 464, and 30 foci in Q1–Q4 respectively). Female neurons only. Aged animals show a progressive shift toward higher *Xist* signal intensity, with no young-animal foci detected in the highest-intensity bin (Q4). **d,** Scatterplot showing the relationship between H3K27me3 intensity within *Xist* foci and total *Xist* intensity for all detected *Xist* objects. Each point represents a single *Xist* focus, colored by age group. A linear regression model was fitted across all pooled samples to assess correlation (slope = 0.4409, adjusted *P* = 2.43 x 10^-6^) **e**, Representative confocal images of NeuN, *Xist*, and pSer5-Pol II in coronal sections from female mice. Merged images show spatial relationship between *Xist* foci and pSer5-Pol II signal. Scale bar, 5 µm. **f**, Quantification of pSer5-Pol II intensity within *Xist* foci across ages in neurons from female mice across age. Each point represents the mean per animal, with error bars indicating mean ± SEM. Statistical analysis was performed using unpaired t-test. n = 1,407 neurons from 15 animals. **g**, Scatterplot showing the relationship between pSer5-Pol II intensity within *Xist* foci and total *Xist* intensity for all detected *Xist* objects. Each point represents a single *Xist* focus, colored by age group. A linear regression model was fitted to assess the global correlation (slope = 0.1750, *P* = 0.008). *Xist* foci are largely excluded from sites of active transcription.

Because *Xist* foci mark the inactive X, we next asked where the age-related H3K27me3 increase is deposited: concentrated on the inactive X, or distributed across the nucleus. To do so, we measured total H3K27me3 signal intensity both across the whole nucleus and within *Xist* foci. Consistent with our western blot data, total H3K27me3 levels within the nucleus of neurons trended up with age (Extended Data Fig. 5d), and consistent with our ChIP-seq data, aged neurons showed a significant increase in H3K27me3 signal intensity within the *Xist* territory compared to young neurons (Fig 5b; p = 0.044). Across neurons, H3K27me3 was preferentially enriched within *Xist* foci relative to the surrounding nucleus, identifying these foci as locally maintained heterochromatin domains on the inactive X.

Bulk intensity averages could obscure whether the *Xist* increase reflects a uniform shift or a growing subset of bright foci, so we next examined its distribution across neurons. We assigned segmented *Xist* puncta (n = 1,986 foci from 18 female animals) to equal-width intensity bins spanning the global *Xist* fluorescence range and compared their distribution by age (chi-square test, X² = 115.03, df = 6, p = 1.80×10⁻²²). Dim puncta (Q1) were distributed roughly equally across age groups (young: 36.0%, middle-aged: 36.0%, aged: 28.1%), whereas brighter puncta were increasingly dominated by aged animals: aged neurons contributed 51.3% of Q3 puncta (versus 33.6% middle-aged and 15.1% young) and 66.7% of the brightest Q4 foci, with no young foci detected in Q4 (Fig 5c). Thus, the age-related *Xist* increase manifests as a growing subpopulation of high intensity foci rather than a uniform brightening across neurons. This intensity gradient in turn allowed us to test whether H3K27me3 scales with *Xist*. Within *Xist* puncta, background-subtracted H3K27me3 enrichment correlated positively with *Xist* intensity (Fig 5d; linear mixed-effects model with animal as a random intercept and age group as a fixed effect; slope = 0.4409, adjusted P = 2.43×10^-06^), indicating that the two marks accumulate together on the inactive X, and that the H3K27me3 increase within *Xist* foci during aging is at least partly distinct from the global upward trend of H3K27me3 with age.

If these *Xist* foci correspond to the inactive X, they should be depleted of active transcription. To test this, we quantified phosphorylated serine 5 of the RNA Polymerase II large subunit C-terminal domain (pSer5-Pol II), which marks transcription initiation, in female hypothalamic neurons (Fig 5e). Overall pSer5-Pol II intensity did not change with age in female neurons, either across the nucleus (Extended Data Fig. 5e), or within *Xist* foci (Figure 5f). Across foci, pSer5-Pol II increased only slightly with *Xist* intensity (Fig 5g; slope = 0.175), a much shallower relationship than that between H3K27me3 and *Xist* (Fig. 5d; slope = 0.44). Together, these data indicate that *Xist* foci, which mark the inactive X, selectively accumulate H3K27me3 but not active RNA Polymerase II with age in female neurons. This localization suggests that the transcriptional and accessibility increases we observed on the female X do not arise from reactivation of the inactive X, but more likely from the active X.

## Discussion

The hypothalamus coordinates with other brain regions and the periphery to regulate many homeostatic processes, including circadian rhythms, appetite control, hormone release, thermoregulation, and memory^64–66^. This region is also altered with age^67^, and therefore has the potential to underlie a number of systemic aging phenotypes. The hypothalamus also mediates some of the longevity benefits of dietary restriction^68^ and is implicated in the regulation of lifespan^69^. Our previous work identified transcriptional changes on the X chromosome, particularly increased *Xist* expression, as a signature of female hypothalamic aging^13^. In this work, we therefore aimed to elucidate the molecular and epigenetic changes associated with hypothalamic aging, and observed extensive sex differences across several regulatory layers including transcription, chromatin accessibility, and histone methylation (H3K27me3).

Our study allowed us to identify specific cell types that are enriched for sex differences during aging. Immune cells stood out as one such cell type, with increases in resident immune cell activation being primarily female-dominated. This manifested as divergent changes between sexes in microglial reactivity markers during aging, with female microglia showing a broad DAM transcript upregulation with age that was not observed in males, which strengthened noticeably at the 24-month old timepoint. The consequences and drivers of these microglial alterations remain unclear, including whether this female-specific DAM upregulation is harmful or adaptive, and if epigenetic or sex-chromosome mechanisms may contribute to these sex specific DAM trajectories.

Our inclusion of a middle-aged (12-month) timepoint proved valuable for interpreting the tempo of hypothalamic aging, not only its endpoint. Aging studies that contrast young and aged animals alone capture only the changes accumulated across life, and cannot resolve when a change arises suddenly or whether it progresses steadily. Here, many of the pronounced female-biased and chrX-enriched transcriptional and accessibility changes, including the microglial shift above, were already established by middle age, indicating that female hypothalamic aging begins earlier than a comparison of young and aged animals alone would reveal. Resolving this tempo has practical consequences: it distinguishes changes that plateau by middle age from those that continue to intensify into old age, and it marks middle age as a candidate window in which sex-specific aging trajectories begin to diverge. More broadly, these observations argue that denser age sampling will be valuable for defining the trajectory, rather than merely the presence, of molecular aging phenotypes.

Deciphering the upstream causes of the sex specific aging changes discussed above and observed in other studies remains a challenge for the field, though our findings suggest a possible involvement of the X chromosome. Sex chromosome content has been linked to sex differences in lifespan^70^ and age-related defects in sex chromosome dosage compensation (e.g., X chromosome inactivation in mammals) has been hypothesized to partially underlie sex-specific aging^71^. In line with this, recent evidence suggests that mammalian X chromosome inactivation is not fully maintained during aging, but rather is subject to age-related de-repression, with evidence emerging in both mice and humans^22–24^. Here, we identified a female-specific upregulation of transcription and chromatin accessibility on the X chromosome with age across multiple cell types and, consistent with our previous study, observed increased *Xist* expression as a female-specific signature of aging in the mouse hypothalamus. Additionally, we found that the *Xist* locus undergoes a progressive increase in chromatin accessibility with age, implicating upstream epigenetic mechanisms for its increased expression with age.

Interestingly, upregulation of *XIST* on the inactive X with age was recently identified in an allele specific analysis of human bulk RNA-seq data^24^, suggesting a possible conservation of this signature across mouse and human aging. Our analysis also re-identified X-linked aging signatures reported by other studies. For example, a recent study in the hippocampus identified age-related escape of *Plp1* as a prominent change^23^; female-specific upregulation of *Plp1* was also present in our dataset, indicating that some X-linked changes may span multiple brain regions during aging.

X chromosome inactivation (XCI) in female mammals involves the deposition and removal of multiple repressive and permissive chromatin marks, respectively^19,21,32,72^, and several of these epigenetic layers (e.g., histone deacetylation and DNA methylation) act synergistically to maintain the repressive state of the inactive X^73^.

Importantly, epigenetic changes are also recognized as a hallmark of aging, and chromatin modifying enzymes (such as histone methyltransferases and deacetylases) can be modulated to extend or shorten the lifespan of model organisms^27–29,74^. XCI’s reliance on extensive epigenomic modifications for its establishment and maintenance opens the possibility that age-related epigenetic changes and attrition of XCI maintenance may be connected.

One modification enriched on the inactive female X chromosome is trimethylation of lysine 27 on histone H3 (H3K27me3), which has been reported to undergo age-related alterations in several species^55–59^. Strikingly, we observed genome-wide redistribution of H3K27me3 and an increase in its total levels with age in the hypothalamus. Through ChIP-seq, we observed that aged hypothalami acquired H3K27me3 autosomal “age domains” – reminiscent of those reported in liver by Yang et al^34^ – over pre-existing low-accessibility heterochromatin and gene-poor regions, indicating that the hypothalamus undergoes genome-wide heterochromatin remodeling with age, possibly marking a transition to a more H3K27me3-reliant form of heterochromatin. These domains also somewhat resemble the H3K27me3 “mesas” previously reported in human senescent cells^75^, and such H3K27me3 domains have also been observed in aged mouse hippocampus^76^. These prior studies observed loss of H3K9me3 at these regions with age and have suggested that the coincident increase of H3K27me3 is a compensatory change^34,76^, and similar H3K27me3 compensation has been seen upon knockout of H3K9-methyltransferases^77^. A recent re-analysis of spike-in normalized single-cell RNA-seq data (from the Tabula Muris Senis atlas^78^) identified a reduction in absolute spike-in normalized transcript abundance in aged mice in several cell types, including neurons and oligodendrocytes^79^, consistent with the global H3K27me3 increase with age we observed here, though we did not observe a global decrease in nuclear pSer5-PolII sum signal intensity with age, arguing against a global repression of transcription during neuronal aging.

Intriguingly, we found that H3K27me3 elevation and redistribution were particularly prominent on the X chromosome in the aging female hypothalamus, but not in males. We validated this increase with H3K27me3 immunohistochemistry and *Xist* co-staining with RNA-FISH, and were able to putatively attribute an age-related increase in H3K27me3 to the inactive X chromosome. Whether the female-specific changes we observed in X-linked transcription, accessibility, and H3K27me3 abundance occur in other brain regions with age remains unknown, although we previously observed increased *Xist* levels with age in the hippocampus^13^. Importantly, what we observed was somewhat discordant: transcription and accessibility increases on the female X chromosome were associated with an increase of repressive markers (*Xist* and H3K27me3). One possibility may be that the latter are compensatory changes in response to a broader dysregulation of heterochromatin on the inactive X. Alternatively, a reinforcement of repression on the inactive X may occur in parallel with age-related hyperactivation of the active X. Consistent with this, we observed that Ser5P-PolII intensity within *Xist* territories did not change significantly with age, which suggests that much of the overall RNA- and ATAC-level upregulation we observed on the female X may originate from the active X. In alignment with this model, recent work reported that in vivo knockout of Sirtuin 7 (*Sirt7*) resulted in disrupted dosage compensation and increased H3K27me3 levels on the inactive X, whereas the active X displayed increased transcription and H3K36ac levels^80^. Importantly, this model is not incompatible with the recent observation that some escape genes increase in expression with age on the inactive X in the hippocampus^23^, given that escape genes are a relatively small number of genes in the mouse (<5%).

The more uniform, flattened H3K27me3 landscape we observed on the aged female X chromosome by ChIP-seq is suggestive of an acquisition of a less ordered XCI state. Though the inactive X is known to be highly enriched for H3K27me3, in future work it would be valuable to assess changes to other XCI-related epigenetic features with age (e.g., H3K27ac, H3K4me1/3, H2AK119Ub, H3K9me2/3, macroH2A, 5mC), and expand these analyses to incorporate allele specific chromatin profiling. Future studies using hybrid mice with polymorphic chromosomes, with one X chromosome copy harboring a deletion in *Xist*, would allow for allele specific bulk profiling of histone modifications on the inactive X chromosome during tissue aging, and may confer insight into age-related alterations to heterochromatin more broadly. Assessing these changes in both sexes will also allow for the improved understanding of the molecular features of hypothalamic aging and their possible effects on sex-specific aging, and how sex chromosome content influences the aging process more generally.

## Methods

### Single-nucleus isolation

Young (3-month-old), middle-aged (12-month-old), and aged (24-month-old) C57BL/6 male and female mice were obtained from the National Institute on Aging. Mice were housed and handled according to protocols approved by the Brown University Institutional Animal Care and Use Committee (IACUC) and in accordance with institutional and national guidelines. Animals were housed at 70 ± 2 °F and 50–70% humidity, and had ad libitum access to Lab Diet 5010 chow and water. Animals were maintained on a 12 h light/12 h dark cycle, with lights on from 07:00 to 19:00. To synchronize the estrous cycle, female mice were exposed to male bedding for three days before euthanasia. All animals were euthanized at Zeitgeber time (ZT) 2–3.

Fresh mouse hypothalamus tissue was dissociated in 500 uL of lysis buffer per sample in a 2 mL Dounce homogenizer. Lysates were combined with 900 uL cold 1.8M sucrose cushion and then gently streamed onto 500 uL cold 1.8M sucrose cushion. Contents were centrifuged at 4 °C for 45 minutes at 13000 rpm. Supernatant was discarded and the pellet was resuspended in 500 uL wash buffer. Contents were spun at 4 °C for 5 minutes at 500 g. Supernatant was discarded and nuclei pellet was resuspended in 250-500uL Nuclei Buffer. All buffers were made fresh on the day of isolation and stored at 4 °C.

| Lysis buffer | Final concentration | Manufacturer and catalog # |
| --- | --- | --- |
| Trizma-HCl pH 7.4 | 10 mM | Sigma T2194 |
| NaCl | 10 mM | Sigma 59222C |
| MgCl<sub>2</sub> | 3 mM | Sigma M1028 |
| Tween-20 | 0.1% | Bio-Rad 1662404 |
| Nonidet P40 Substitute | 0.1% | Millipore 492018 |
| Digitonin | 0.005% | Thermo Fisher BN2006 |
| BSA | 1% | Miltenyi Biotec<br>130-091-376 |
| DTT | 1 mM | Sigma 646563 |
| RNAse Inhibitor | 1 U/uL | Sigma 3335402001 |
| Nuclease Free Water |  |  |
| **Wash buffer** |  |  |
| Trizma-HCl pH 7.4 | 10 mM | Sigma T2194 |
| NaCl | 10 mM | Sigma 59222C |
| MgCl<sub>2</sub> | 3 mM | Sigma M1028 |
| Tween-20 | 0.1% | Bio-Rad 1662404 |
| BSA | 1% | Miltenyi Biotec<br>130-091-376 |
| DTT | 1 mM | Sigma 646563 |
| RNAse Inhibitor | 1 U/uL | Sigma 3335402001 |
| Nuclease Free Water |  |  |
| **1.8M sucrose cushion** |  | Sigma NUC201-1KT |
| 2M sucrose cushion solution | 90% | Sigma S99308 |
| Sucrose cushion buffer | 10% | Sigma S9058 |
| **Nuclei buffer** |  |  |
| Nuclei buffer (20X) | 1X | 10x Genomics<br>2000153/2000207 |
| DTT | 1 mM | Sigma 646563 |
| RNAse Inhibitor | 1 U/uL | Sigma 3335402001 |
| Nuclease free water |  |  |

### Statistics and reproducibility

No statistical methods were used to predetermine sample sizes, but our sample sizes are similar to those reported in a previous study using single-cell RNA-seq on brain tissue^81^. All statistical tests were two-sided unless otherwise stated. No animals were excluded from the study. Individual nuclei were excluded from analysis based on quality control metrics (feature count and mitochondrial read count). Data collection and analysis of the sequencing experiments were not performed blind to the conditions of the experiments.

### snMultiome: data collection, quality control, data processing, and analysis

snRNAseq and snATACseq libraries were constructed according to the 10x Genomics Chromium Next GEM Single Cell Multiome ATAC and Gene Expression protocol (CG000338 Rev E). A target of 8,050 nuclei were loaded onto the Chromium Next GEM Chip J (with the exception of 1YF, for which 7,375 nuclei were loaded). GE samples were assessed for quality using a High Sensitivity DNA BioAnalyzer assay, and ATAC samples were assessed for quality using a Fragment Analyzer. Sequencing was performed by Azenta Life Sciences using the Illumina NovaSeq.

Reads were aligned to the mm10 (2020) mouse reference genome using Cell Ranger ARC v2.0.1. Downstream data processing and analysis were conducted in R v4.4.0, using ArchR v1.0.2^43^ and Signac v1.13.0^82^ for snATAC-seq analysis and Seurat v5.1.0^83^ for snRNA-seq analysis. ATAC-seq quality control (QC) was performed following the ArchR tutorial with default parameters, and doublets were removed. Peaks were called on pseudobulk aggregates using MACS2^44^, combining fragments from major cell types to improve signal resolution. Differential expression (DE) analysis was performed in Seurat using the FindMarkers function with the Wilcoxon rank-sum test (test.use = “wilcox”).

### Imaging and analysis of H3K27me3 and Pol II within *Xist* foci *in vivo*

Young (4 month), middle (12 month), and aged (22-24 month) C57BL/6 female and male mice were obtained from the National Institute of Aging. Aged females (22 months) and aged males (24 months) were combined into a single aged cohort for analysis given equivalent proportions of age-related pathology at these timepoints. N = 6 females and 6 males per age group. Animals were group-housed whenever possible. Mice were housed and used according to protocols approved by Buck Institute IACUC and in accordance with institutional and national guidelines. Animals were housed at 70 +/- 2 °F with humidity from 50-70%. Animals were fed ad libitum Envigo Teklad 18% Protein chow and water. The light cycle was 12 h on/12 h off; lights were on between 6:00 and 18:00.

### Perfusion, sectioning

Mice were anesthetized with Avertin (125-300 mg/kg IP) and perfused with heparin/PBS followed by 4% paraformaldehyde. Brains were removed and post-fixed in 4% paraformaldehyde overnight at 4 °C and dehydrated with a 30% sucrose gradient. After 2-3 days, the brains were embedded in Tissue-Tek® O.C.T. Compound and stored at −80 °C. All brains were cut into 40 *μ*m coronal sections using a cryostat (Leica CM1950 Cryostat) and stored in a cryoprotectant buffer (30% ethylene glycol, 30% glycerol in 0.05 M phosphate buffer) at -20 °C until use and within 6 months.

### Staining and hybridization

Sections were blocked in 10% Normal Donkey Serum (Jackson ImmunoResearch) and 1% Triton X-100 in 1X phosphate buffered saline (pH 7.4) for one hour at room temperature, then incubated with a mouse monoclonal anti-NeuN primary antibody (1:500; Millipore Sigma; MAB377 clone A60) and a rabbit polyclonal anti-H3K27me3 primary antibody (1.7 µg/mL; ActiveMotif; Cat. 39155; Lot 2325432-11) in 10% Normal Donkey Serum and 0.1% Triton X-100 overnight at 4 °C. Sections were washed in PBS and 0.01% Triton X-100 and incubated with an Alexa Fluor 488 donkey anti-mouse secondary antibody (1:500; Invitrogen) and Alexa Fluor 647 donkey anti-rabbit secondary antibody (1:500; Invitrogen) for two hours at room temperature. Sections were washed in PBS and 0.01% Triton X-100 and then PBS only. Sections were adhered to SuperFrost slides and dried in the dark.

Following immunofluorescence staining, RNA fluorescence in situ hybridization for *Xist* RNA was performed using LGC Biosearch Technologies Stellaris FISH Probe Mouse *Xist* with Quasar 570 Dye (Biosearch Technologies; VSMF-3094-5), following the manufacturer’s protocol for fresh frozen mouse brain tissue instructions with minor modifications: sections were secured with a hydrophobic barrier pen rather than a coverslip during hybridization; hybridization was extended to 18 hours; post-hybridization Wash Buffer A and Wash Buffer B washes each included additional buffer-exchange steps; DAPI counterstaining concentration was adjusted; an Autofluorescence Eliminator treatment was incorporated into the post-hybridization dehydration series; and Prolong Diamond antifade mountant was substituted for Prolong Gold. Briefly, sections were post-fixed in 3.7% formaldehyde for 15 minutes, dehydrated through an ethanol series, and hybridized with fluorescently labeled *Xist* probes at 37 °C for 18 hours. Post-hybridization washes were performed under high-stringency conditions to reduce non-specific signals. Sections were counterstained with DAPI (1 µg/mL) in Wash Buffer A solution for 30 minutes, washed in Wash Buffer B, and washed in 1X PBS. Sections were then dehydrated through an ethanol series that incorporated a 5-minute treatment with Autofluorescence Eliminator (Millipore Sigma cat. 2160) to reduce lipofuscin-associated background. Sections were mounted with Prolong Diamond antifade mountant and cover glass, cured overnight at room temperature in the dark, and subsequently stored at 4 °C. All staining conditions, imaging parameters, and probe concentrations were kept constant across experimental groups. Staining for H3K27me3/NeuN and for PolII/NeuN were performed on separate adjacent sections from the same animals.

### Imaging

Brain sections were imaged on a Zeiss LSM700 Confocal Microscope equipped with a 63x objective (NA 1.4) for quantification analysis. Image bit depth is 16-bit. Z-stacks were collected at 0.56 µm intervals using identical laser power, detector gain, and pinhole settings for all experimental conditions. Maximum intensity projections were used for analysis. Investigators were blinded to the identity of each sample while staining, imaging, and segmentation. Each brain section was imaged twice, once on each side of the third ventricle to capture the Arcuate Nucleus in the hypothalamus.

### Nuclear and neuronal segmentation

Segmentation was performed using Zeiss Zen Blue software (Version 3.11) via the ‘Segment Region Classes Independently’ processing method. The DAPI signal was used to segment all nuclei using an intensity-based thresholding and watershed algorithm. Each DAPI-positive nucleus was assigned a unique object identifier. NeuN staining was used to classify these nuclei as neuronal or non-neuronal. Specifically, the NeuN signal was evaluated within each DAPI-defined nuclear region, and nuclei exceeding a predefined intensity threshold were automatically classified as NeuN-positive neurons.

Segmentation thresholds were empirically determined and held constant across all images and conditions (intensity ranges: DAPI = 9172-65535; NeuN = 11592-37642; H3K27me3 = 5640-59828; PolII = 3524-61802; *Xist* = 15017-45004). These thresholds were manually validated on a subset of images to confirm accurate detection of total number of cells, neurons, H3K27me3- or Pol II-positive neurons, and *Xist* foci). For NeuN-positive nuclei, H3K27me3 signal was quantified within the corresponding NeuN-defined nuclear mask. Mean fluorescence intensity, area, and integrated density were extracted for each nucleus. *Xist* RNA puncta were detected within NeuN-defined nuclear regions using spot detection based on size and pixel intensity criteria (size range = 23000-36150, intensity threshold = 482514-4933475). Multiple *Xist* puncta could be assigned to a single nucleus. For each nucleus, the total number of puncta, sum puncta intensity, and total puncta area were recorded.

### Data aggregation and statistical analysis

Quantitative measurements (area, mean intensity, sum intensity) from Zeiss ZEN Blue software (version 3.11) were exported as CSV files and processed in R (version 4.4.2 (2024-10-31)) using the tidyverse collection of packages^84^ (dplyr, tidyr, readr, stringr, purrr, ggplot2). Per-nucleus and per-punctum measurements were retained for all analyses. For summary statistics, measurements were aggregated hierarchically: individual foci or nuclei were averaged within each image, images were averaged within each animal, and each animal was treated as a biological replicate. Aggregate data were exported to Microsoft Excel using the writexl package and imported into GraphPad Prism (version 11) for visualization and group comparisons. Statistical comparisons between age groups were performed using Welch’s unpaired t-test with Bonferroni correction for three pairwise comparisons (young vs. middle, young vs. aged, middle vs. aged), as indicated in figure legends. The distribution of *Xist* puncta across fluorescence intensity bins was compared across age groups using the chi-square test of independence (base R, R version 4.4.2).

The relationship between *Xist* signal intensity and marker enrichment within *Xist* foci was visualized using ordinary least squares (OLS) linear regression (base R lm()). Statistical inference was performed using a linear mixed effects model (lme4 and lmerTest packages) with animal identity as a random intercept to account for the non-independence of multiple *Xist* foci measured per animal. Age group was included as a fixed effect. Model comparisons were performed using likelihood ratio tests to assess whether age moderated the slope of the *Xist*-marker relationship. Robust regression (MASS::rlm, MM-estimator) was used as a sensitivity analysis to confirm that results were not driven by high-intensity outlier foci.

*Xist* RNA FISH puncta from H3K27me3 (n = 949 foci, 18 animals) and PolII (n = 1,037 foci, 15 animals) imaging datasets were pooled for analysis of *Xist* intensity distribution across age. Both datasets imaged the same *Xist* channel (T2) under identical acquisition conditions. Unpaired comparison of *Xist* intensity for the 15 animals present in both datasets showed no systematic difference between imaging sessions (Wilcoxon signed-rank, W = 76.0, p = 0.379). Overall distribution overlap was 79.9% (KS D = 0.173). The 22-month age group showed greater inter-dataset variability (KS D = 0.469, overlap 55.6%), reflecting high within-cohort biological variability in aged animals rather than a systematic batch effect.

### Trajectory Analysis

Trajectory (or “pseudotime”) analysis is a commonly used method where a trajectory is fit to an embedding space, after which signatures (often transcription or chromatin accessibility) across this trajectory can be assessed to study a continuum of states across single cells to model processes such as development and differentiation. To further characterize sex differences in our snMultiome dataset’s immune cell population, we performed trajectory analysis across the immune subset on our transcriptional UMAP embedding using the Monocle3 package^51^. We chose to map this trajectory as a single joint trajectory across all immune cells for both sexes to be on a comparable pseudotime axis. Prior to trajectory mapping we filtered out 11 of 2,456 total immune cells with outlier UMAP coordinates to avoid these cells biasing the trajectory. We programmatically calculated a root principal node using Monocle3’s “get_earliest_principal_node” function, setting the young condition as the “time_bin” parameter. Per-gene accessibility was quantified as gene body accessibility extended 2kb upstream of the transcription start site (TSS) using the “GeneActivity” function from the Signac package^82^ in R. GO Biological Process over-representation analyses were performed with clusterProfiler^85^, using ENTREZ IDS derived from gene symbols, where resulting terms with Benjamini-Hochberg adjusted p < 0.05 were retained for plotting.

### ChIP-seq

ChIP experiments were performed using the SimpleChIP® Plus Sonication Chromatin IP Kit (#56383, Cell Signaling Technology), following manufacturer instructions and scaling reagent volumes down to accommodate the lower average starting tissue mass of individual whole hypothalami relative to other commonly profiled organs/regions. Flash frozen hypothalami were individually minced into small pieces and fixed in fresh methanol-free formaldehyde (#12606, Cell Signaling Technology) at a final concentration of 1% for 10 minutes. The Covaris M220 system was used to sonicate each sample for 8 minutes (5% duty factor, 200 cycles/burst, 75 peak incident power). Each chromatin sample’s concentration was estimated by reverse crosslinking a small aliquot from each sample and measuring its DNA concentration with the Qubit HS dsDNA Quantitation kit (Q32851). Chromatin samples were equalized by concentration and volume before proceeding to avoid potential biases in initial tissue mass. For spike-in normalization, a small, equal quantity of *Drosophila* spike-in chromatin (Active Motif Cat No. 53083) was added to each chromatin sample prior to splitting initial chromatin samples into ChIP-designated and 10%-input fractions. All ChIP reactions were performed with anti-H3K27me3 primary antibody overnight at 4C (#9733, Cell Signaling Technology). ChIP DNA was quantified using the Qubit HS dsDNA Quantitation kit (Q32851). All ChIP-seq animals were dissected together as one batch and experimentally processed together as a single batch. Sequencing libraries were prepared, multiplexed, and sequenced by Azenta Life Sciences using the Illumina NovaSeq XPlus instrument to generate paired end 150bp reads.

### Western blot

Flash frozen hypothalamus samples were minced and suspended in a hypotonic lysis buffer (10mM Tris-HCl pH 8, 1mM KCl, 1.5mM MgCl2, 10mM Sodium Butyrate, 1mM DTT, 0.2% IGEPAL CA630, 1 cOmplete™ Mini EDTA-free Protease Inhibitor Cocktail, buffer adapted from Shechter et al.^86^) and dounce homogenized with 15 strokes of the “loose” pestle and 15 strokes of the “tight” pestle. Nuclei were washed after an 800xg spin at 4C, which was then followed by an additional 800xg spin and resuspension in 0.2M HCl for overnight histone extraction at 4C, followed by adjustment to neutral pH and quantification of supernatant with Qubit protein assay (Q33211). For the 3-month-old and 21-month-old comparisons, histone extracts were mixed with Laemmli buffer and loaded by equal mass on a precast gradient gel for SDS-page, followed by transfer to a 0.2 µm nitrocellulose membrane. After transfer, membrane was cut in half, using one side for probing H3K27me3 (Cell Signaling Technology, #9733), and the other for probing histone H3 (Cell Signaling, #4620). For the western blot comparing 3-month-old and 24-month-old samples, we first probed for H3K27me3, followed by stripping and re-probing for histone H3 (Restore Western Blot Stripping Buffer Cat. No. #21059). Blocking was performed in 5% Milk TBS for histone H3, and 5% BSA TBS for H3K27me3, followed by overnight incubation at 4C in 5% BSA TBS with primary antibody. Membranes were then washed with TBST and incubated with a HRP-conjugated secondary antibody. Next, membranes were washed again with TBST and developed in Pierce ECL western blotting substrate, followed by chemiluminescent detection with the ChemiDoc MP imaging system.

### ChIP-seq data processing

Raw FASTQ reads were trimmed in paired end mode using fastp v0.23.2^87^ with default parameters. Next, paired-end, trimmed FASTQ files were aligned to the mouse*-Drosophila* (mm10-dm6) concatenated genome using bowtie2 v2.4.1^88^ with the no-mixed, no-discordant, and ‘-X 1000’ parameters. Aligned BAM files were then filtered for deduplicated, properly paired, primary reads with MAPQ >=30 using samtools v1.13^89^, followed by calculation of spike-in mapping rates. Next, *Drosophila*-mapping reads were filtered out from these BAMs, followed by removal of reads overlapping Encyclopedia of DNA Elements (ENCODE) mm10 blacklisted regions^90^ for downstream analysis and visualization of mouse reads.

*Drosophila* fragment counts from the primary, properly paired, MAPQ filtered, deduplicated BAM files were obtained with the “featureCounts” command from subread v2.1.1^91^. These *Drosophila* fragments were then pre-calibrated to account for initial spike-in chromatin loading (see below section in Methods). DESeq2 v1.46.0^62^ was then used to estimate size factors from these pre-calibrated drosophila counts. Mouse fragment counts for the primary DESeq2 analysis were obtained using subread’s featureCounts command on the primary, properly paired, MAPQ filtered, deduplicated BAM files with *Drosophila* reads filtered out and ENCODE blacklist regions removed. Primary DESeq2 analyses used combined models (“∼ sex + age”) and within-sex age models (“∼ age”). A set of DEseq2 results were obtained using spike-in normalization, and an additional set was obtained using standard, default DESeq2 normalization.

For calling broad H3K27me3 domains, we used Enriched Domain Detector (EDD) v1.1.19^92^ with the “required fraction of informative bins” parameter set to 0.97 and an FDR threshold of 0.05. EDD calls were performed on pooled aged ChIP reads using their matched pooled aged input reads as controls. Genome browser tracks were visualized using pyGenometracks v3.9^93^.

### ChIP-seq analysis

Spike-in normalization was applied in two scenarios: (1) creating scaled BigWigs for downstream visualization of genome tracks; and (2) for differential enrichment analysis. To create spike-in normalized BigWig files, the following formula^94^ was used to calculate a multiplicative scale-factor for a given sample i:

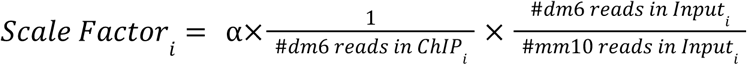

For each sample, this scale factor was then applied to the sample’s ChIP BAM file using the bamCoverage command from deeptools v3.5.6^95^. Correcting for variability in the initial addition of spike-in chromatin (estimated by the spike-in mapping rate in the sample’s respective input), minimizes technical noise in scale-factors that is frequently observed in spike-in normalization^96^. For input-normalized ChIP genome browser tracks, for each sample, ChIP and matched input coverage were CPM-normalized in 20bp genomic bins and combined as the per-bin log2 ratio, log2[(CPM_ChIP + 1) / (CPM_input +1)], using the deepTools bamCompare command. Principal component analysis was performed on input-normalized BigWigs generated using the same procedure, except with 5kb bins.

To incorporate spike-in information into differential enrichment analysis, ChIP read counts were passed into DESeq2 along with custom size factors derived from *Drosophila* ChIP read counts, as has been previously described^94,97^. Following these descriptions, to correct for variability in initial loading of spike-in chromatin, *Drosophila* read counts in each ChIP sample were pre-calibrated to account for technical variability in addition of spike-in chromatin to samples prior to immunoprecipitation. The following formula was used to obtain a pre-calibration factor for each animal i:

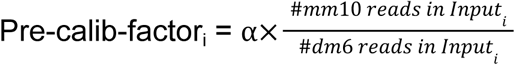

where α is a coefficient applied to all samples such that the largest pre-calibration factor is 1. For example, an animal with a higher proportion of dm6 reads initially present in the input would imply an inflated spike-in mapping rate in the ChIP sample, which would then be corrected downward by the pre-calibration factor. Each animal’s pre-calibration factor was applied to the *Drosophila* fragment counts from that animal’s ChIP sample, and size factors were then estimated by DESeq2. These size factors were then passed into DESeq2 along with mouse ChIP fragment counts to assess differential enrichment between young and aged ChIP samples.

To test whether age-associated changes in H3K27me3 at the X-inactivation center were extreme relative to the rest of the X chromosome, we employed a two-sided equal-width genomic window permutation test. Let *T*(*W*) denote the mean spike-in-normalized Old-versus-young log_2_ fold-change across 5-kb bins overlapping genomic window *W*. The observed statistic was *Tobs* = *T*(*XIC*), where XIC coordinates were chrX:103184059-103981285. We generated an empirical null distribution by sampling *B* = 10, 000 same-width (as the XIC) windows randomly positioned along the X chromosome (but not overlapping the XIC) with replacement and computing *Ti* for each sampled window. Two-sided extremeness was evaluated after centering on the empirical mean of the sampled windows, 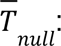 b was the number of sampled windows satisfying 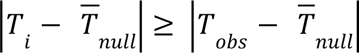, with a resulting empirical p-value of (*b* + 1)/(*B* + 1).

## Acknowledgements

The authors thank members of the Webb lab for critical feedback on the study.

## Author contributions

A.E.W. conceptualized, designed, and supervised the study. L.H. and K.H.H. performed the snMultiome experiments. D.Y., I.O., L.H., and K.H.H. processed and analyzed the snMultiome data. I.O. performed and analyzed the ChIP-seq and western blot experiments. N.A.S. performed and analyzed the staining and imaging of epigenomic marks. D.Y., I.O., N.A.S., L.H., and A.E.W. wrote and edited the manuscript. H.M. reviewed the manuscript and provided feedback.

---

**Extended Data 1.**
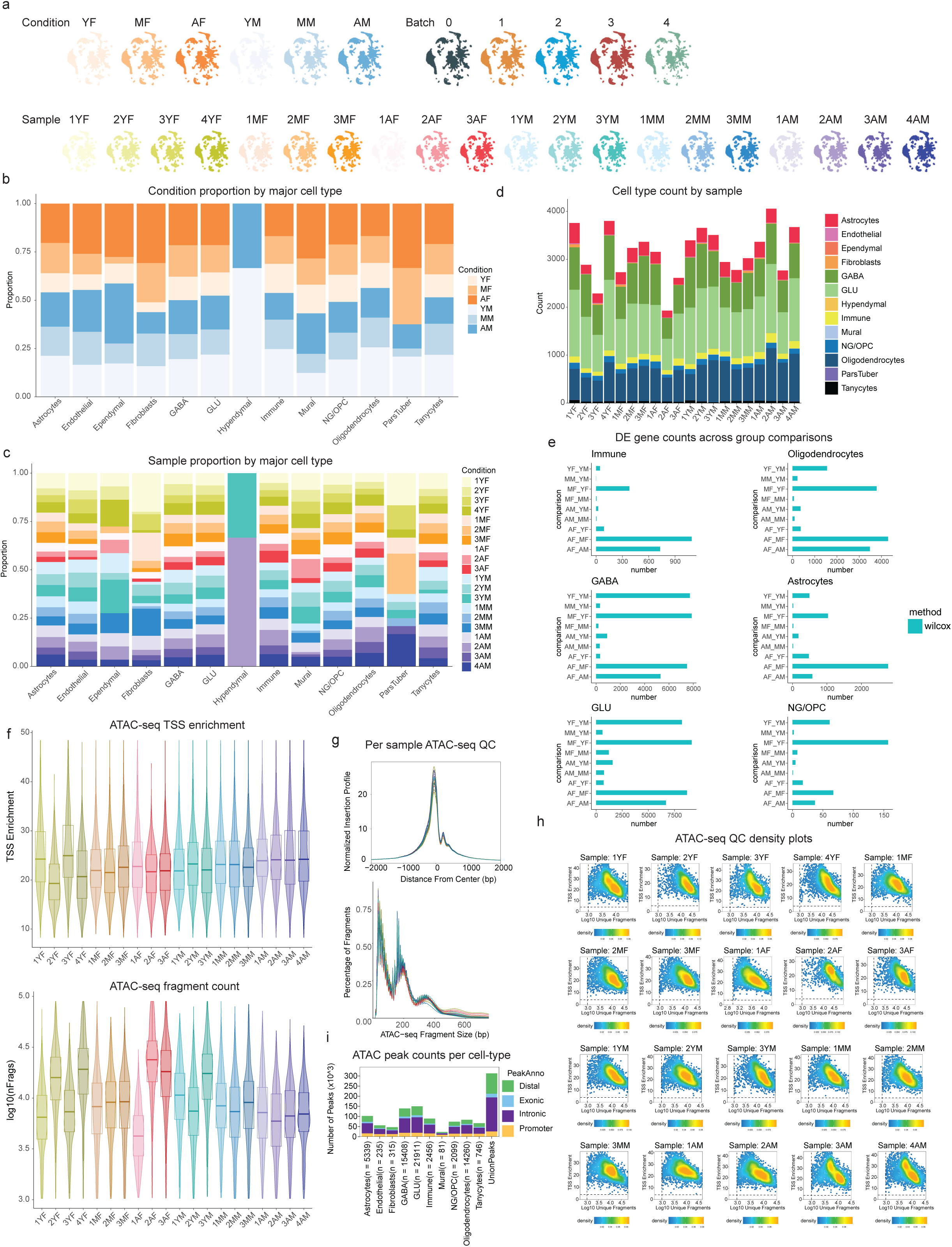
Quality control and cell type distribution in aging hypothalamus multi-omics dataset. **a**, UMAP visualization of nuclei colored by condition, batch, and sample, showing the integration of snMulti-omics data. **b**, Stacked bar plot showing the sex and age groups in individual major cell types. **c**, Stacked bar plot showing experimental animals/samples in individual major cell types. **d**, Bar plot showing the number of nuclei per cell type for each sample. **e**, Cell-type-specific differentially expressed gene (DEG) comparisons across conditions: Bar plots showing the number of significant DEGs in major cell types analyzed using the Wilcoxon test. **f**, Violin plots of TSS enrichment and log10(fragment counts) across samples. **g**, TSS enrichment profiles and fragment size distributions by sample. **h**, Density plot of TSS enrichment vs log10(fragment counts) across samples. **i**, Peak annotation across major cell types.

**Extended Data Figure 2.**
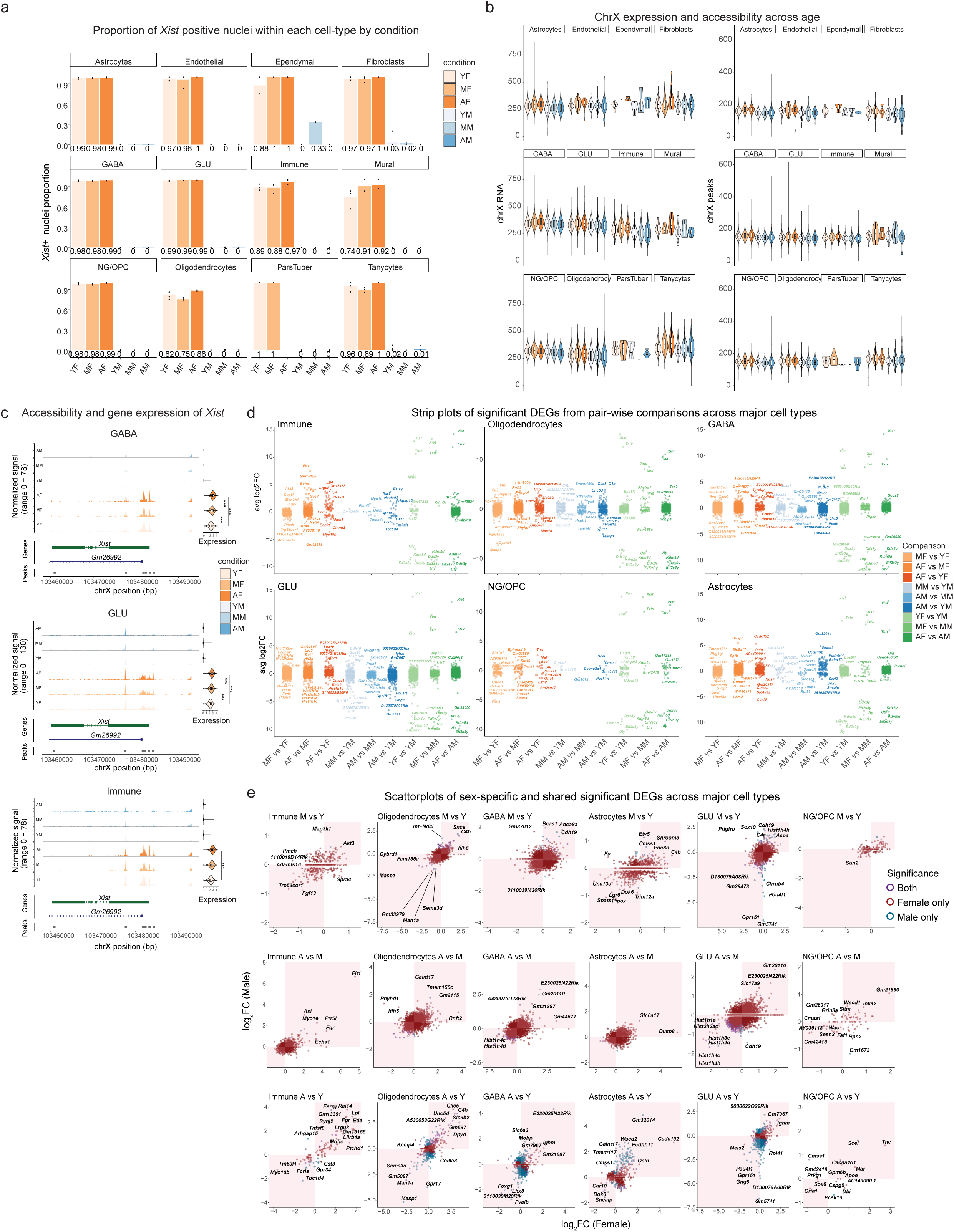
Sex-specific gene expression and chromatin accessibility changes in major cell types with age. **a**, Bar plots showing the proportion of *Xist*-positive cells across major hypothalamic cell types, stratified by sex and age. **b**, Violin plots of chrX gene expression and chromatin accessibility across conditions in major cell types. **c**, Aggregated peaks and expression of *Xist* in major cell types. Statistical significance for expression: Bonferroni-adjusted p-values from the Wilcoxon test; *adjusted p < 0.05, **adjusted p < 0.01, ***adjusted p < 0.001. **d**, Strip plots of significant DEGs from pairwise comparisons across major cell types, highlighting sex- and age-specific expression patterns. **e**, Scatterplots of sex-specific and shared significant DEGs across major cell types.

**Extended Data Figure 3.**
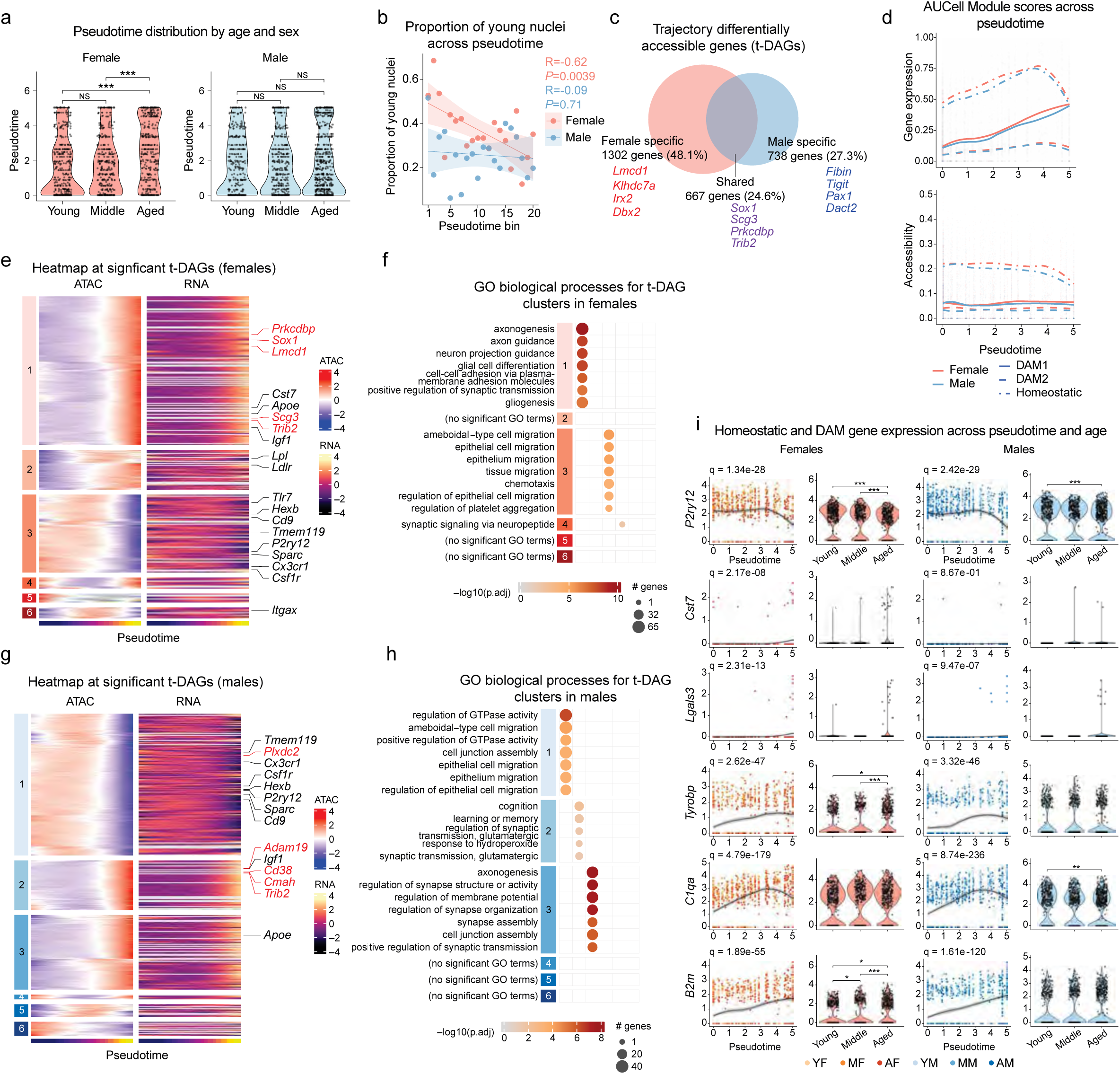
Analysis of accessibility dynamics across pseudotime in male and female microglia. **a,** Violin plots of pseudotime versus age in both sexes, where each dot represents a single nucleus. Statistics: two-sided, unpaired Wilcoxon rank-sum tests on individual nuclei, with three pairwise comparisons per sex; p-values were Holm-Bonferroni adjusted across all six comparisons (***p.adj <0.001) (from left to right: p.adj = 0.10, 1.35×10^-10^, 2.79×10^-6^, 1, 1, 1). **b,** Overlaid scatterplots showing, within each sex, the proportion of nuclei within each quantile bin that are young. Pearson correlations are shown. **c,** Comparison of female versus male t-DA genes across the trajectory identified within each sex by Moran’s *I* test (Benjamini-Hochberg adjusted q-value (FDR) < 0.05). Accessibility for each gene was measured across the gene body and extended 2 kb upstream of the transcription start site. Top 4 significant DE genes within each slice are listed. **d,** AUCell module scores of DAM1, DAM2, and homeostatic microglial signatures in immune nuclei showing aggregated expression (top) and aggregated accessibility (bottom) across pseudotime. **e,** Heatmap of female t-DA genes across pseudotime in female immune nuclei; genes labeled in red are the top five most significant by Moran’s *I* test. **f,** GO BP term summary of t-DA gene clusters from (e). Plotted are top terms with Benjamini-Hochberg adjusted p < 0.05. **g,** Heatmap of male t-DA genes across pseudotime in male immune nuclei; genes labeled in red are the top five most significant by Moran’s *I* test. **h,** GO BP term summary of t-DA gene clusters from (g). Plotted are top terms with Benjamini-Hochberg adjusted p < 0.05. **i,** Normalized expression of additional individual homeostatic and DAM marker genes across pseudotime and age. Expression across nuclei is fitted by a LOESS curve (black line).

**Extended Data Figure 4.**
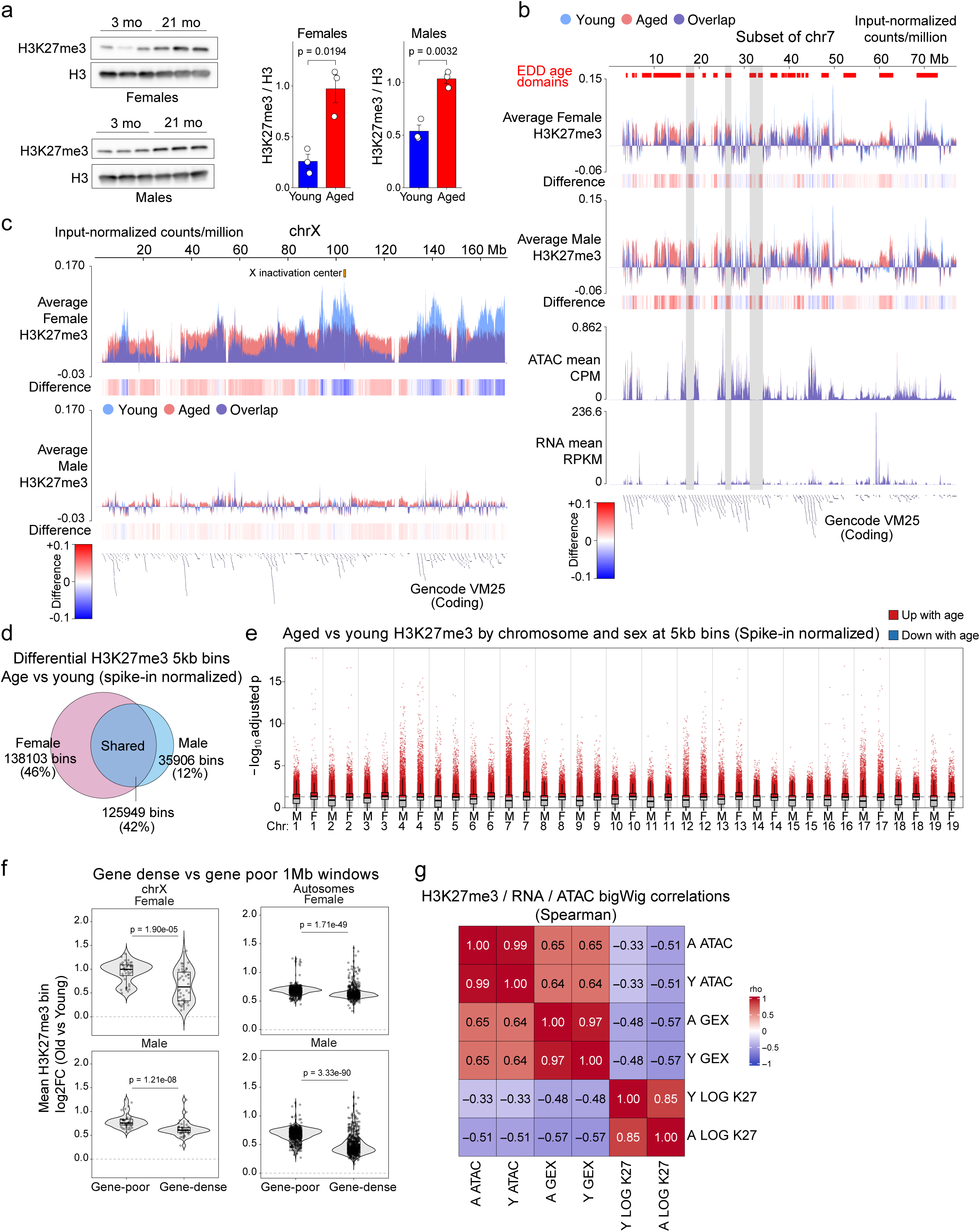
Characterization of age-associated H3K27me3 increases and integration with snRNA-seq and snATAC-seq data. **a,** H3K27me3 western blot of acid-extracted histones from hypothalamus samples of 3- and 21-month-old mice of both sexes (left). Western blot quantifications (right). P-values were determined by Welch’s t-test (n=3 biological replicates per age group). **b,** Genome browser tracks of input-normalized log2cpm H3K27me3 signal, across a region of chromosome 7. H3K27me3 enriched domain detector (EDD) domain calls are shown by red bars above the tracks. “ATAC mean” tracks show the average snMultiome ATAC-seq CPM signal with aged (red) and young (blue) tracks overlaid. “RNA mean” tracks show the averaged snMultiome RPKM (reads per kilobase million) signal for young and aged. Grey shaded bars highlight examples of age domains tightly coinciding with lowly accessible, transcriptionally inactive regions. **c,** X chromosome genome browser tracks of input-normalized log2cpm H3K27me3 signal. **d,** Comparison of female versus male differentially bound 5-kb bins identified by H3K27me3 DESeq2 analysis (spike-in normalized). DESeq2 was run independently within females and within males using an identical universe of bins; bins with |log2FC| >= 0.5 and p.adj < 0.05 were classified as differentially bound. **e,** Between-sex comparison of spike-in normalized H3K27me3 DESeq2 results for 5-kb bins across all autosomes. Significantly upregulated bins (log2FC >= 0.5 and p.adj < 0.05) are colored red and non-significant bins are colored grey. **f,** Comparison of spike-in normalized H3K27me3 log_2_ fold changes across non-overlapping 1-Mb windows in gene-dense and gene-poor regions (protein-coding genes). Within chrX, windows were ranked according to the number of overlapping protein-coding genes (GENCODE vM25 annotation) and grouped into quartiles, with the same ranking and grouping procedure being applied across all autosomes. **g,** Spearman correlations between gene expression, accessibility, and H3K27me3 profiles. Individual snMultiome ATAC BigWig files were averaged into one single mean BigWig per age group. The same averaging procedure was applied to the snMultiome gene expression (GEX) BigWig files, and the input-normalized bulk H3K27me3 BigWig files.

**Extended Data Figure 5.**
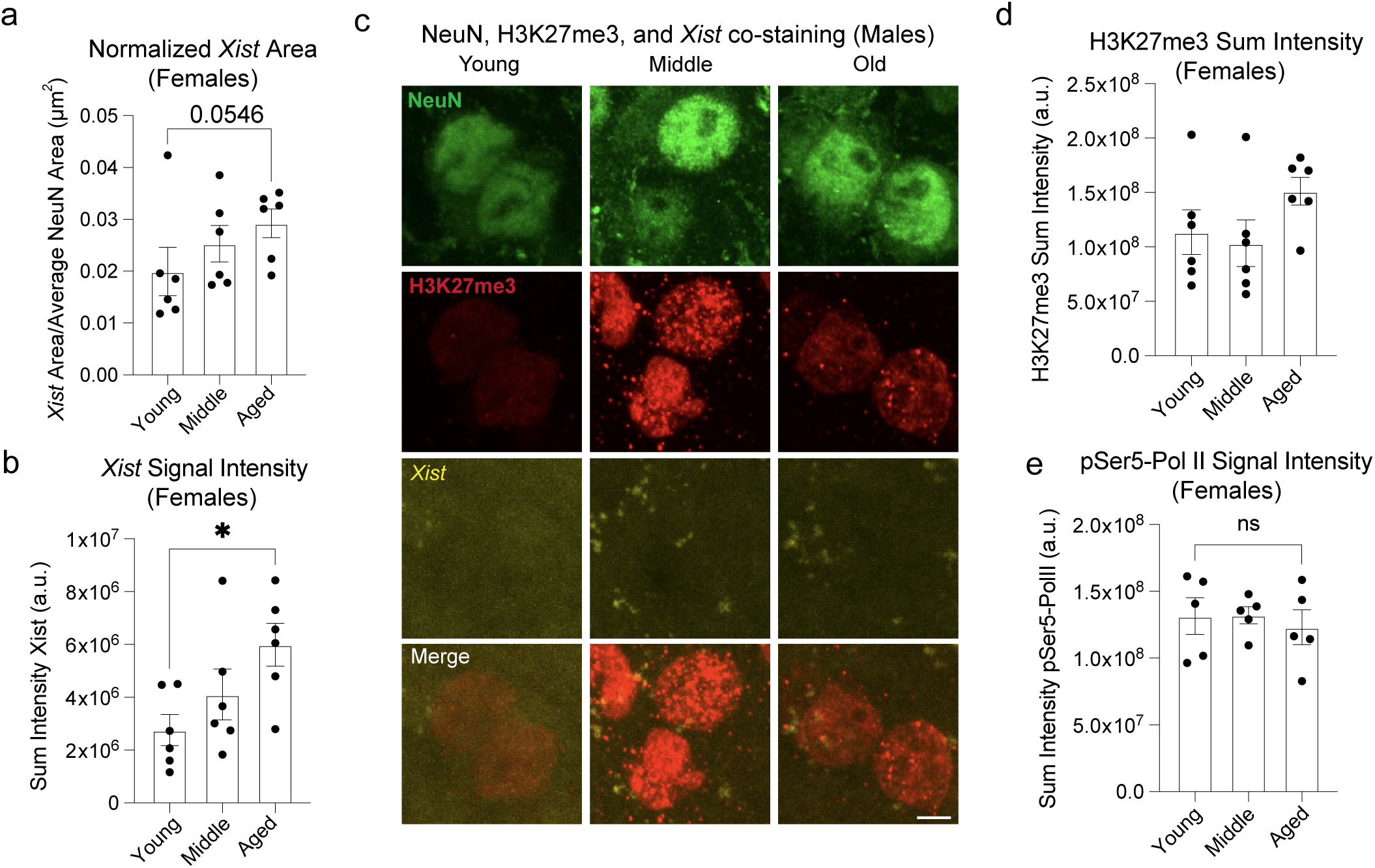
Age-dependent increase in *Xist* intensity in female hypothalamic neurons is not accompanied by changes in pSer5-Pol II levels. **a**, Quantification of *Xist*-positive area per neuron normalized to the average area of DAPI per image across age groups in female mice. Each point represents the mean across all neurons per animal, with bars indicating mean ± SEM. No significant difference was detected using unpaired t-test. **b**, Quantification of *Xist* sum intensity signal per neuron across age groups in female mice. Each point represents the mean across all neurons per animal, with bars indicating mean ± SEM. Statistical significance was assessed using unpaired t-test (p = 0.0101). **c,** Representative confocal images of NeuN, *Xist*, and H3K27me3 in coronal sections of the arcuate nucleus in the hypothalamus from Young (4-month-old), Middle-aged (12-month-old), and Aged (24-month-old) male mice. *Xist* RNA signal is undetectable in male neurons, consistent with the absence of X chromosome inactivation in male somatic cells. Scale bar, 5 µm. **d**, Quantification of H3K27me3 sum intensity within the whole cell across neurons from Young (4-month-old), Middle-aged (12-month-old), and Aged (22-month-old) female mice. Each point represents the mean across all neurons per animal, with bars indicating mean ± SEM. **e**, Quantification of pSer5-Pol II sum intensity within the whole cell across neurons from 4-, 12-, and 22-month-old female mice. Each point represents the mean across all neurons per animal, with bars indicating mean ± SEM. No significant difference was detected using unpaired t-test. n = 1,407 neurons from 15 animals.

**Supplementary Table 1.** Differential expression analysis across major cell types and in pseudobulk across all cells. Wilcoxon rank-sum test results for pairwise age comparisons within each sex; genes with Bonferroni-adjusted p < 0.05 were designated differentially expressed. For the pseudobulk analysis across all cells, DESeq2 was used, with genes at BH-adjusted p < 0.05 designated differentially expressed.

**Supplementary Table 2.** Trajectory-dependent gene expression and accessibility and their associated functional enrichment categories in male and female immune cells.

Complete, unfiltered Moran’s *I* test results for RNA and ATAC modalities along the RNA-derived trajectory, analyzed separately in females and males. Genes with Benjamini-Hochberg-adjusted p values (q values) < 0.05 were designated trajectory differentially expressed genes (tDEGs) or trajectory differentially accessible (tDA genes). Enrichment results were obtained using clusterProfiler, with all genes analyzed in the corresponding trajectory analysis supplied as the background universe.

**Supplementary Table 3.** Age-associated changes in H3K27me3 across genomic bins. DESeq2 results comparing aged versus young mouse H3K27me3 across 5-kb genomic bins. Six worksheets contain results using spike-in or standard normalization, for females, males, or both sexes combined. Tables report genomic coordinates (mm10), log2 fold change, and associated statistics. Positive log2 fold change indicates higher H3K27me3 enrichment in aged samples. Bins were classified as significantly differentially enriched at a BH-adjusted p-value < 0.05 and an absolute log2 fold change > 0.5. Only significant bins were kept for the file size purpose.

